# Oscillatory flow and steady streaming of cerebrospinal fluid in cranial subarachnoid space

**DOI:** 10.64898/2026.03.25.714044

**Authors:** Mariia Dvoriashyna, Jaco J.M. Zwanenburg, Alain Goriely

## Abstract

Cerebrospinal fluid (CSF) is a Newtonian fluid that bathes the brain and spinal cord and oscillates in response to the physiological periodic changes in brain volume, of which the cardiac cycle is a major driver. Understanding this motion is essential for clarifying its contribution to solute transport, waste clearance, and drug delivery. In this work, we study oscillatory and steady streaming flow in the cranial subarachnoid space using a lubrication-based theoretical framework. The model represents the cranial CSF compartment as a thin fluid layer bounded internally by the brain surface and externally by the dura, driven by time-dependent brain surface displacements. We first derive simplified governing equations for flow over an arbitrary smooth sphere-like brain surface and obtain analytical solutions for an idealised spherical geometry with uniform displacements. We then incorporate realistic displacement fields reconstructed from MRI measurements in healthy subjects and solve the reduced equations numerically. The results show that oscillatory forcing produces a steady streaming component that may enhance solute transport compared with diffusion alone. This work provides a mechanistic description of the flow generated by physiological brain motion and highlights the potential presence of steady streaming in cranial subarachnoid fluid dynamics.

## 1 Introduction

Cerebrospinal fluid (CSF) is a Newtonian fluid with physical properties similar to water (Linninger *et al*., 2016), that bathes the brain and the spinal cord. CSF is located within the brain’s ventricular system, in the subarachnoid space (SAS; see figure 1a) of spine and cranium, and in the perivascular spaces (PVS) surrounding the blood vessels (Kelley & Thomas, 2023). The main functions of the CSF flow include regulating intracranial pressure (ICP), transporting nutrients, and removing waste products from brain metabolic activity (Rasmussen *et al*., 2022).

**Figure 1:**
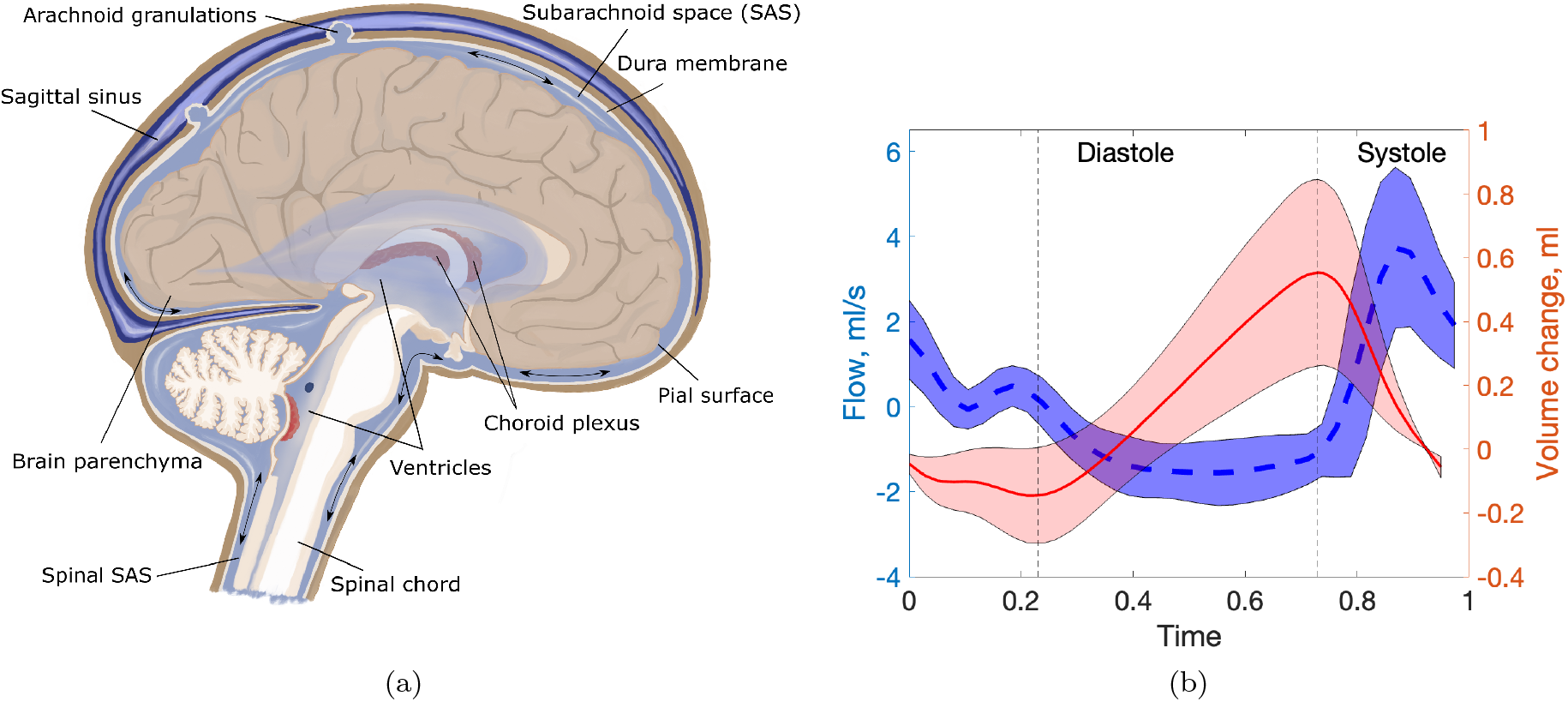
(a) Sketch of the cross-section of the brain. The arrows indicate oscillatory CSF motion. The drawing is based on the image in commons.wikimedia.org. (b) Left axis: flow from the brain into the spine (positive if fluid flows into the spine), measured using PC-MRI (Adams *et al*., 2020). The line represents the mean value over 8 subjects, and the shading is the standard deviation. The *x*-axis is the scaled time by the duration of a heartbeat. Right axis: change of CSF volume during the cardiac cycle, from Adams *et al*. (2020). Vertical dashed lines separate systolic and diastolic phases.

During the cardiac cycle, the CSF undergoes an oscillatory motion, which can be measured in the SAS with phase-contrast magnetic resonance imaging (PC-MRI) (Feinberg *et al*., 1985; de Marco *et al*., 2004; Baledent *et al*., 2006), while recent advances have enabled the quantification of the CSF mobility in the PVS (Hirschler *et al*., 2025). During systole, the inflow of blood into the arteries results in an increase in brain volume. The Monro-Kellie doctrine (Monro, 1783) posits that since the skull is rigid, the combined volume of blood, CSF, and brain should remain constant in time. Thus, as shown in figure 1(b), with arterial expansion during systole, the CSF is expected to move from the cranium to the spine (Linninger *et al*., 2005; Geregele *et al*., 2019). Conversely, during diastole, brain volume decreases and CSF returns to the cranial compartment.

Pulsatile flow of CSF is known to play a crucial role in enhancing solute transport in the PVS (Thomas, 2019; Vinje *et al*., 2021; Bojarskaite *et al*., 2022; Asgari *et al*., 2016) and in the spinal SAS (Lawrence *et al*., 2019; Salerno *et al*., 2020), with implications in solute clearance from the brain. Although the clearance mechanisms are not yet fully understood, all current theories agree on the involvement of CSF flow (Ringstad & Eide, 2024). Moreover, enhancement of solute transport has practical applications in drug delivery through intrathecal drug administration, in which a drug is injected within the spinal canal, specifically inside the subarachnoid space where CSF circulates, and is transported by the CSF to the brain (Tangen *et al*., 2019; De Andres *et al*., 2022; Ayansiji *et al*., 2023). In order to understand mechanisms responsible for solute transport in CSF, we require a detailed description of the fluid flow.

CSF also plays role in ICP regulation (Linninger *et al*., 2016). While temporal variation of ICP can be measured with an invasive procedure where probes are placed into the ventricles or brain parenchyma, or less invasively with lumbar puncture (Evensen & Eide, 2020), these methods cannot comprehensively assess spatial variations of the ICP within or between compartments in the central nervous system (CNS). Mathematical modelling and simulations can quantify spatial profiles of the ICP (Causemann *et al*., 2022).

Analytical models of CSF flow based on lubrication theory were previously developed for spinal SAS (*e*.*g*. Sánchez *et al*., 2018; Cardillo & Camporeale, 2021; Sincomb *et al*., 2022), along with computational works, including realistic geometries (Khani *et al*., 2018; Coenen *et al*., 2019; Gutiérrez-Montes *et al*., 2021) and refined numerical methods (Toro *et al*., 2019, 2022). Interestingly, Sánchez *et al*. (2018) found that inertial steady streaming is generated in spinal SAS, which plays an important role in solute transport. Further studies also addressed the dispersion of solutes in the spine in the context of drug delivery (Gutiérrez-Montes *et al*., 2021; Salerno *et al*., 2020; Lawrence *et al*., 2019; Ayansiji *et al*., 2025). Several works (Hsu *et al*., 2012; Tangen *et al*., 2015, 2017) studied intrathecal drug delivery numerically, accounting for both cranial and spinal CSF compartments.

Here, to complement such studies, we focus on the CSF flow in cranial SAS (cSAS). Indeed, fewer theoretical studies are available in cSAS. Two computational models address CSF flow in cSAS (Gupta *et al*., 2010) and SAS between the spine and cranium (Gupta *et al*., 2009), both using realistic geometry from MRI data. In particular, Gupta *et al*. (2010) used a flux condition corresponding to the change in cSAS volume on a fixed pial boundary to prescribe uniform brain displacements and imposed velocities on the boundary with the spine obtained from PC-MRI data. Further, Sweetman & Linninger (2011) and Sweetman *et al*. (2011) studied a fluid-structure interaction problem for the cranium and the entire CNS, comparing the results with Cine PC-MRI measurements. The authors prescribed the displacements of the ventricular surfaces and of the parts of SAS closest to blood vessels with the highest density. Similarly, Howden *et al*. (2011) used the prescribed velocity of ventricular walls as a driver for the CSF. However, ventricular pumping only accounts for about 10 % CSF volume exchange between the spine and the cranium (Geregele *et al*., 2019). Causemann *et al*. (2022) proposed a computational framework that combines a poroelastic description of the brain parenchyma and all the cranial CSF compartments. These authors accounted for blood inflow using uniformly distributed source/sink terms in the parenchyma and obtained tissue displacement and CSF flow. Further reviews of computational works on CSF flow can be found in Linninger *et al*. (2016), Kurtcuoglu *et al*. (2019) and Kelley & Thomas (2023). However, to date, there has been no detailed study of CSF flow in cSAS, accounting for realistic brain displacements.

In this work, we develop a model of CSF flow in cSAS informed by data-informed displacements of the brain surface, and use it to predict both ICP gradients in cSAS and solute transport. Brain tissue displacement during the cardiac cycle can be measured using Displacement ENcoding via Stimulated Echoes (DENSE) technique (Adams *et al*., 2019, 2020). This is an MRI-based method that can quantify the submillimetric displacements associated with brain tissue motion. These authors report displacement maps and an associated change of CSF volume during the cardiac cycle of around 0.74 mL, comparable with other studies (*e*.*g*. Stoquart-El Sankari *et al*., 2009; Wåhlin *et al*., 2012). Herein, we prescribe brain surface displacements approximated from the data in Adams *et al*. (2020) and assume they are the sole driver of fluid flow in cSAS.

To model fluid transport, we use lubrication theory, an asymptotic technique valid in long and thin domains. This approach complements existing numerical works and provides a detailed understanding of the fluid flow and its dependency on the model parameters. Furthermore, it allows us to compute inertial steady streaming in cSAS, which would be challenging to do using direct numerical simulations due to the thin space and time-dependency of the flow. We first derive lubrication equations for an arbitrary sphere-like surface of the brain, i.e. any surface that can be parametrised by a function *R*(*θ, ϕ*) in spherical coordinates. To build intuition, we derive an analytical solution for a simplified case of uniform displacements and spherical brain in §2.6. We then solve the generic case of a realistic brain with displacements prescribed from brain data from Adams *et al*. (2020), which we approximate with spherical harmonics in §3. Results are presented in §4 and discussed in §5.

## 2 Model set-up and simplification

We assume that the CSF flow in cSAS is driven by the prescribed displacement of brain surface (also referred to as the *pial surface*). Since we extract the displacement fields from in-vivo MRI data, we assume that the brain moves as a rigid solid. In other words, we assume that fluid pressure in the cSAS does not displace the pial surface. For simplicity, here we will neglect the presence of arachnoid trabeculae in cSAS and consider that cSAS is an open space filled with CSF.

### 2.1 Geometry and coordinate system

We assume that the brain has a smooth shape (no gyri or sulci). We place the origin on the same vertical axis as the outlet to the spine. Then, as shown in figure 2, we parameterise the sphere-like surface 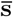 of the brain using a radial function 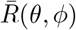 in spherical coordinates

**Figure 2:**
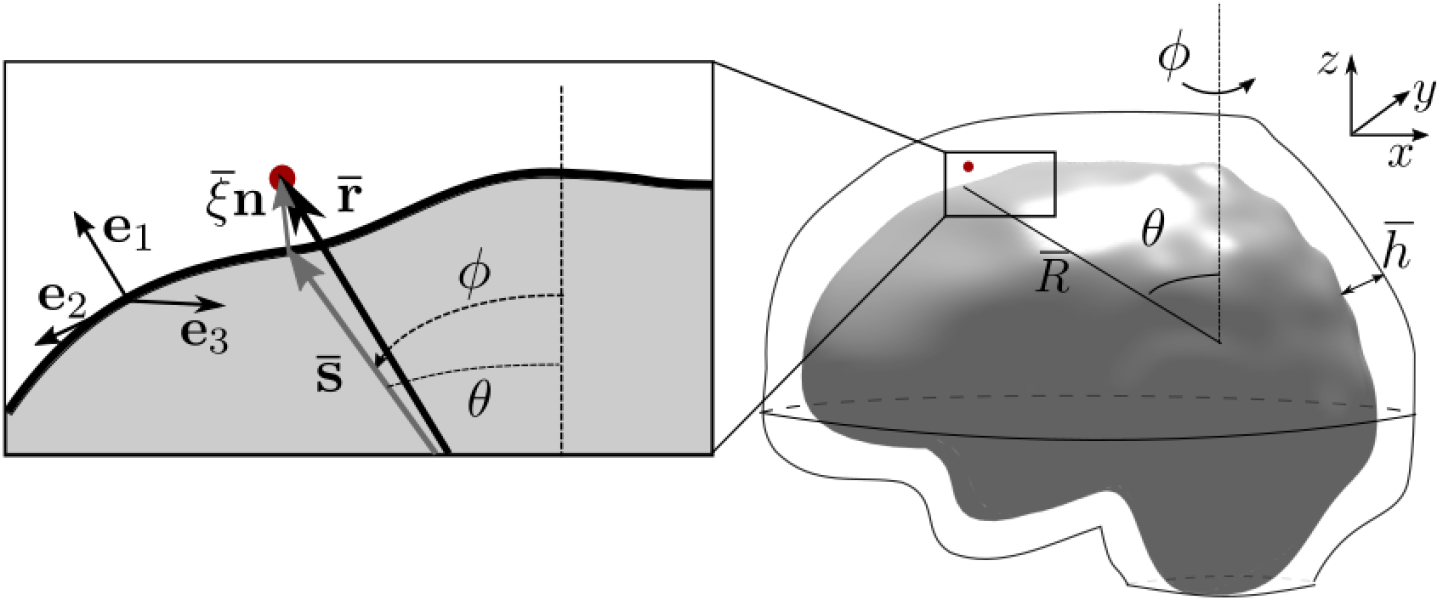
Sketch of the domain and the coordinate system. The spherical-like brain is shown on the right, surrounded by the dura membrane. Right: zenithal and azimuthal coordinates *θ* and *ϕ* and spatially variable radius of the brain 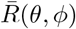. The thickness of SAS is 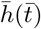. Left: zoom-in to the curved surface of the brain. The position vector 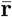 for the red point is shown as a sum of vector 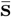 and 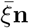 (2). (**e**_1_, **e**_2_, **e**_3_) are base vectors on the brain surface.

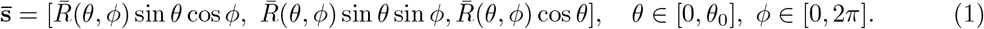

We choose 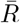 to be either constant (spherical approximation) or a distance from the origin to the realistic brain surface described in §3.1. For a fixed pial boundary, the cSAS is assumed to have a uniform thickness *h*_0_. Using values from table 1, we observe that there is a separation of length scales between the thickness and the length of the cSAS, with aspect ratio *ϵ* = *h*_0_*/R*_0_ ≈ 0.027, where *R*_0_ is a typical length for the brain radius. Thus, assuming further that the inverse of the curvature of the surface 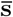 is always much larger than the SAS thickness, we can write the position vector 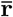 of a point in the cSAS as

**Table 1:** Model parameters and their values.

| Parameter | Value | Source |
| --- | --- | --- |
| $R_0$, characteristic radius of the brain | $7.5 \cdot 10^{-2}$ m | §3.3 |
| $h_0$, characteristic thickness of cSAS | $2 \cdot 10^{-3}$ m | 0.5-3 mm (Frydrychowski *et al.*, 2012) |
| $\theta_0$, zenithal span of the domain | 3 | Estimated from spinal SAS diameter of 17 mm (Ulbrich *et al.*, 2014) |
| $\omega/2\pi$, heart rate | 1 Hz | |
| $\mu$, CSF viscosity | $0.8 \cdot 10^{-3}$ Pa s | Causemann *et al.* (2022) |
| $\rho$, CSF density | 1007 kg/m<sup>3</sup> | Barber *et al.* (1970) |

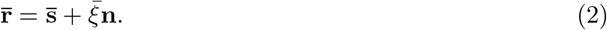

where 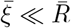. We thus adopt the coordinate system (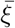, *θ, ϕ*) to describe material points in the cSAS (see figure 2).

As usual, it is convenient to work in terms of dimensionless variables, and we scale all lengths w.r.t *R*_0_ so that 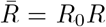. The local basis vectors are set to be two (dimensionless) tangent vectors to the pial surface and a unit normal vector,

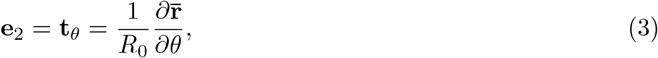

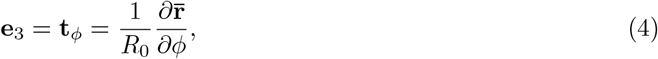

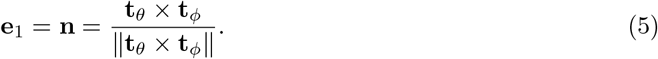

These vectors are given explicitly in (43). Note that this basis is not orthogonal, as 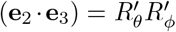, where we denote 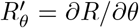, and 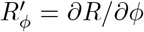. Neglecting the 𝒪 (*ϵ*) terms, the metric tensor for this basis is

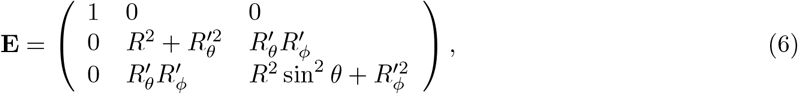

where **E**_*ij*_ = *e*_*ij*_ = (**e**_*i*_ · **e**_*j*_). We denote the determinant of this matrix as 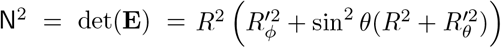. More details on this coordinate system and the conjugate metric tensor are given in Appendix A.

We assume that the surface of the brain undergoes periodic displacements with frequency *ω*. Owing to the small aspect ratio *ϵ* of the domain, the displacement in the normal **e**_1_ direction will result in flows 1*/ϵ* larger than those due to the displacements in the tangential directions. Thus, we assume that the brain surface only moves in the **e**_1_ direction with magnitude 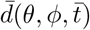. The cSAS spans between 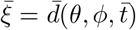 and a prescribed cSAS thickness at rest 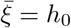.

### 2.2 Governing equations

The CSF is modelled as an incompressible Newtonian fluid with viscosity *μ* and density *ρ* (see table 1). The CSF motion can then be described by the Navier-Stokes equations for an incompressible fluid, which read (with overbars denoting dimensional variables):

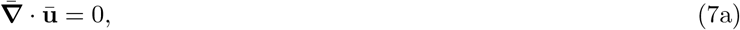

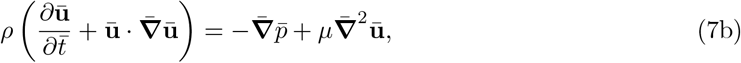

where 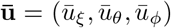 is the velocity vector and 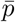 is the pressure. We impose a no-slip boundary condition on the pial surface 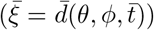 and at the dura membrane 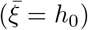. Furthermore, since the spine serves as an outlet for the fluid from cSAS, we impose zero pressure at the boundary with the spinal SAS *θ* = *θ*_0_.

### 2.3 Model simplification

The characteristic thickness of cSAS is approximately *h*_0_ ≈ 2 mm (Frydrychowski *et al*., 2012). From the experimental observations shown in Fig. 1b, the mean peak change of CSF volume during each oscillation is ΔV = max (*V*) −min (*V*) ≈0.74 ml (Adams *et al*., 2020). If the brain displaces uniformly in the direction normal to its surface, the change of the volume in SAS is approximately proportional to the change in its thickness, and scales as ΔV ∼ Δh = max(*h*) − min(*h*). Assuming a spherical brain with radius *R*_0_ from table 1, the displacement of the pial surface is about 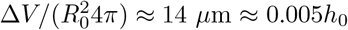. The real brain displacements are spatially non-uniform, and those in normal direction reach values of 35 *μ*m≈ 0.0175*h*_0_ at the equatorial plane (Adams *et al*., 2020). Therefore, in this work, we assume that Δ*h/h*_0_ ∼ 𝒪 (*ϵ*).

Since the domain is long and thin with aspect ratio *ϵ* ≪ 1, we use lubrication theory to simplify the problem. It is convenient to work in terms of dimensionless variables, and we introduce the following scalings

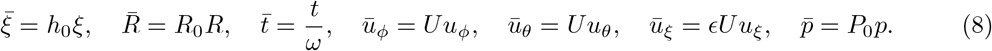

We pick the velocity scale *U* based on the frequency of oscillations, *U* = *ωh*_0_ ≈ 12.6 cm/s. A convenient scale for the pressure is *P*_0_ = *ω*^2^*h*_0_*Rρ*, which balances the pressure term with the leading acceleration term. Note that with this scaling the derivative in the normal direction scales as 1*/h*_0_, rather than 1*/R*_0_, so that 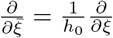.

Using this scaling to non-dimensionalise the equations (7), we obtain two dimensionless groups: the aspect ratio *ϵ* and the Womersley number 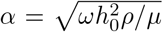. Using parameters from table 1, we estimate them to be

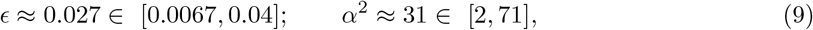

where the ranges in brackets are obtained by varying the value of *h*_0_ ∈ [0.5, 3] mm. Following standard lubrication assumptions, we neglect the contributions of order *ϵ* ^2^ and *ϵ* ^2^*/α*^2^ and smaller. The resulting momentum equations in cSAS (7b) read (see Appendix A for the full derivation)

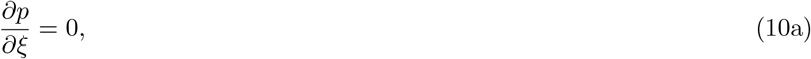

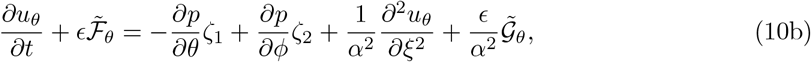

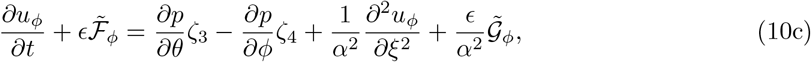

where the functions *ζ*_*i*_ depend on *R* and its first derivatives and are reported in (55). The non-linear terms 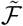 are as follows

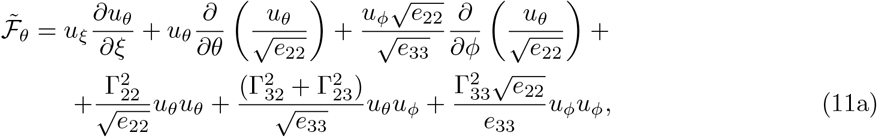

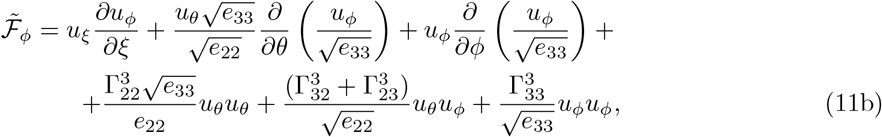

where 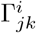 are Christoffel symbols (74). The terms 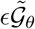, and 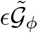 are the remaining linear terms of order *ϵ*. Note that (10a) implies that pressure is independent of *ξ* and thus *p* = *p*(*θ, ϕ, t*).

The continuity equation (7a) in our coordinate system is

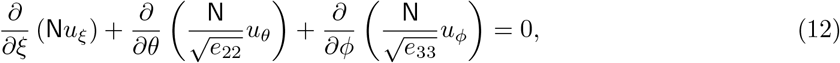

where N = ∥**t**_θ_ × **t**_*ϕ*_∥ is given explicitly in (50). It is simpler to work in terms of the fixed domain rather than the moving one, so we rescale the transversal coordinate, 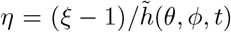, where 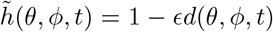. Here 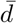 is scaled with *h*_0_*ϵ*, so that the resulting dimensionless function *d* is of order 1. The new coordinate *η* has a fixed range, *η* ∈ [−1, 0]. The system of equations in the fixed domain reads

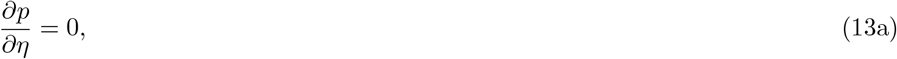

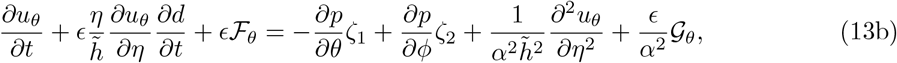

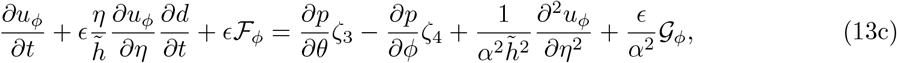

and the continuity equation is

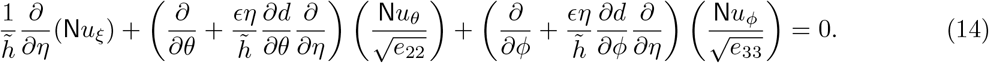

The functions ℱ and 𝒢 are the inertial terms and residual linear terms in the new coordinates (see (57)).

At the pial boundary (*η* = −1), we prescribe the velocity **v** = (*v*_*ξ*_, 0, 0), so that, **u**(−1, θ, *ϕ, t*) = **v**. At the dura membrane (*η* = 0), we have the no-slip boundary condition, **u**(0, *θ, ϕ, t*) = **0** and at *θ* = *θ*_0_, *p* = 0.

We now return to the governing equations (13) and expand the solution in powers of *ϵ*

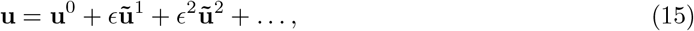

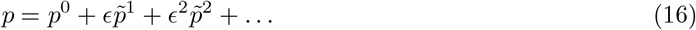

In the next sections, we will consider separately the leading order flow and the steady streaming flow to first order. We note that even though the leading viscous terms, which have factors 1*/α*^2^, are of order *ϵ* for the values in table 1, we choose to treat them as terms of 𝒪 (1), as they appear as 𝒪 (1) in the boundary layers. A similar approach was used by Blondeaux & Vittori (1994), who showed that it leads to the same result as a formal boundary layer analysis.

### 2.4 Oscillatory flow at O(1)

To leading order, (13) becomes a system of linear PDEs

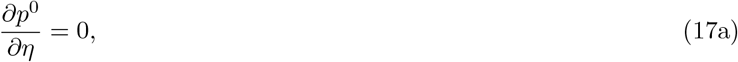

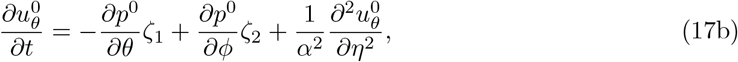

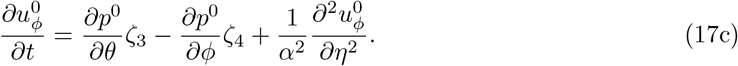

The first equation implies that *p*^0^ = *p*^0^(*θ, ϕ, t*). We expand the solution of this system in *N*_*h*_ time harmonics as follows

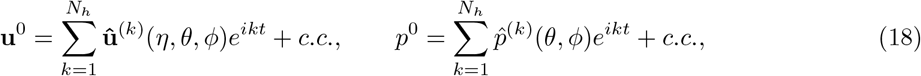

which leads, for each harmonic *k*, equations for the amplitudes

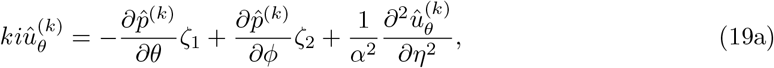

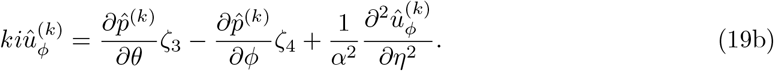

Since 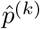 does not depend on *η*, we can solve the equations (19) and apply no-slip boundary conditions for 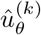 and 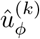 to obtain

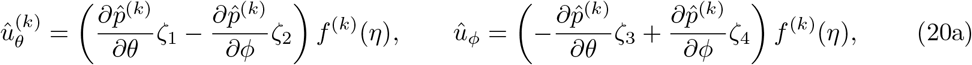

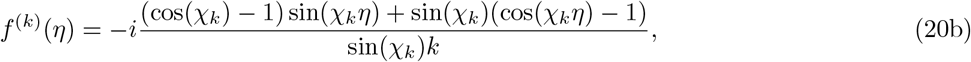

where 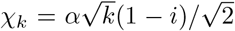. Using these expressions, we integrate the continuity equation (14) to leading order with respect to *η* ∈ [−1, 0] and apply boundary conditions 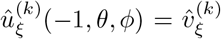 and 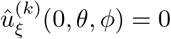 to obtain the equation for 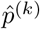:

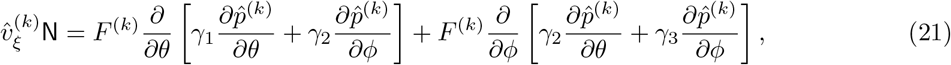

where

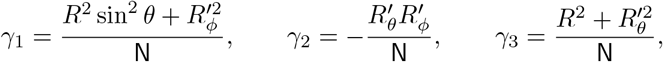

and 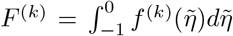. Therefore, we have the governing equation for the pressure to leading order with the boundary condition

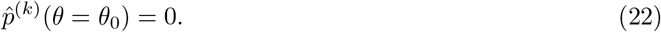

Finally, the third component of the velocity, 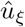 can be found by integrating the continuity equation:

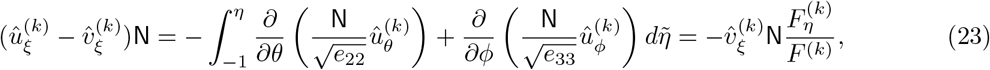

where for the last equality we used (20), (21) and defined

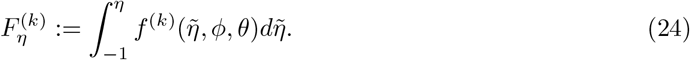

Thus, the velocity in the normal direction is 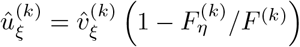, where the dependency on *θ* and *ϕ* comes only from the factor 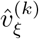.

### 2.5 Steady streaming 𝒪 (*ϵ*)

We now consider steady solution to (13), (14) at order *ϵ*, which we denote with **u**^1^, *p*^1^. This is 0-th harmonic of the Fourier series expansion of **ũ**^1^, 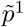 (the 𝒪 (*ϵ*) solution). The momentum equations (10) for **u**^1^, *p*^1^ are as follows

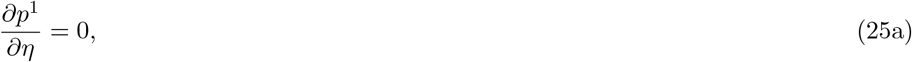

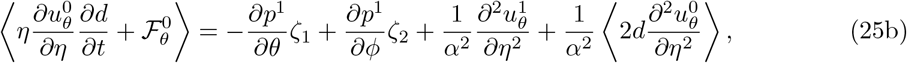

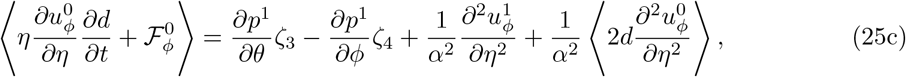

where 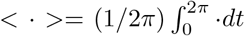 denotes the average over a period of oscillations. The inertial terms 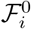 are obtained from substituting the leading-order solution into (57) and keeping the leading-order terms. The first-order continuity equation for the steady flow reads then:

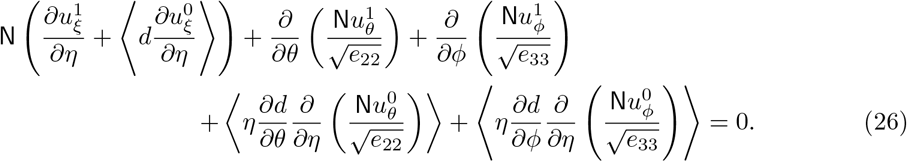

Noting that *p*^1^ = *p*^1^(*θ, ϕ, t*), we integrate equation (25b) twice and apply no-slip boundary conditions to obtain

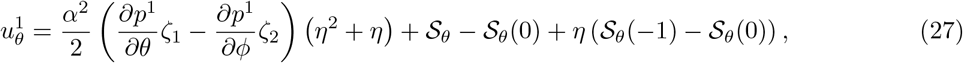

where

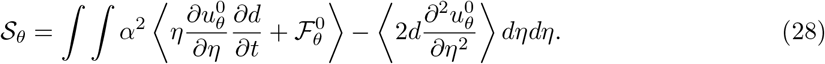

The analogous expression can be obtained for 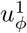. Finally, we integrate the first-order continuity equation (26) to obtain the equation for the steady streaming pressure *p*^1^, which reads

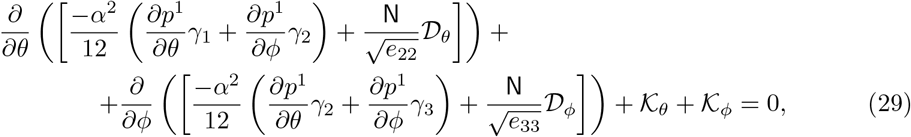

where

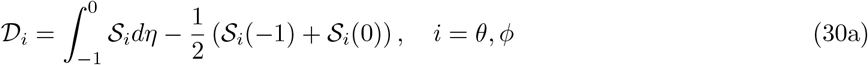

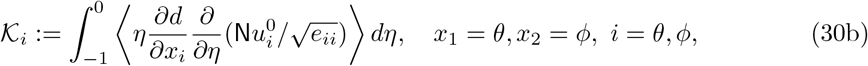

(no summation on repeated indices). To obtain the above expression we used 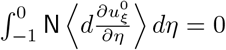 (see (65)). After solving (29) for the pressure we can find the tangential velocities from (27) (and analogous one for 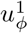) and normal velocity from integrating (26).

### 2.6 Analytical solution for spherical brain

#### 2.6.1 Leading order

In a simplified case of a spherical brain (*R* = 1) and spatially uniform brain displacements 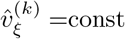, an analytical solution can be found. It is axisymmetric with respect to *ϕ* and the equation for the pressure reads

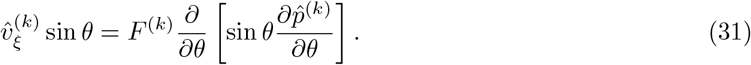

where *F* ^(*k*)^ is a constant. Integrating twice and applying the boundary conditions on *p*, we obtain

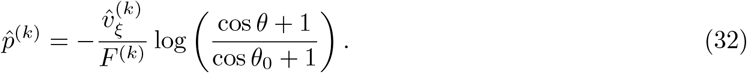

The velocity components are obtained from (20) and (23):

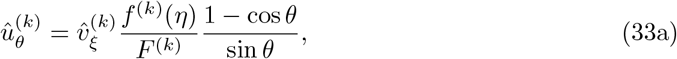

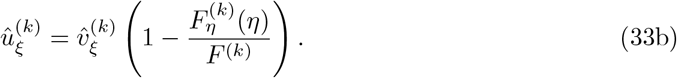

#### 2.6.2 Steady streaming

The *θ*-momentum equation for the steady streaming reads

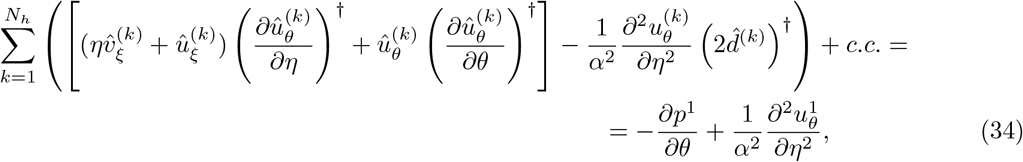

where the superscript † denotes complex conjugate. We will use the analytical solution for the leading order to write the forcing term as

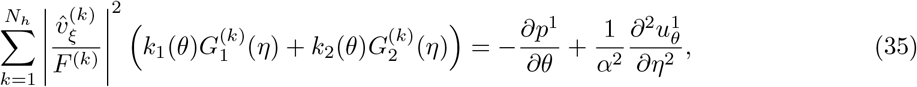

where we introduced the following notations

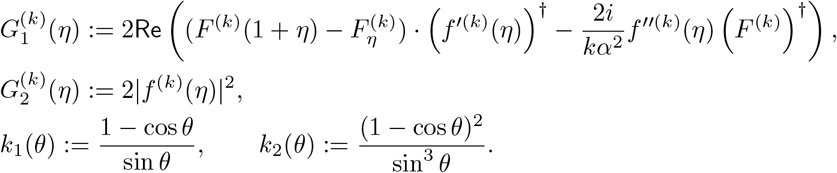

Integrating twice and applying the boundary conditions, we obtain (similar to (27))

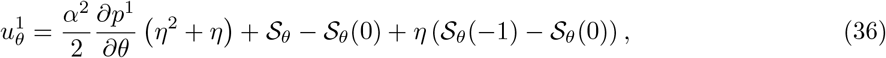

with

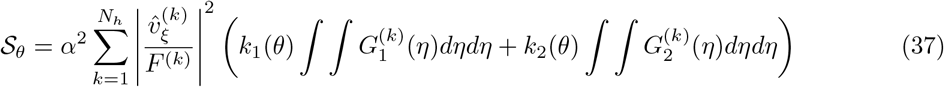

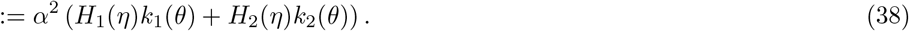

The equation for pressure (29) reads

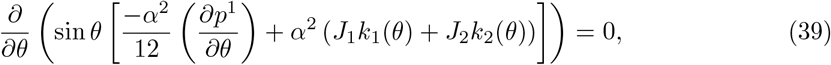

where we used 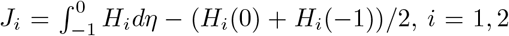. Integrating this equation twice and imposing zero flux at *θ* = 0 and zero pressure at *θ* = *θ*_0_, we obtain

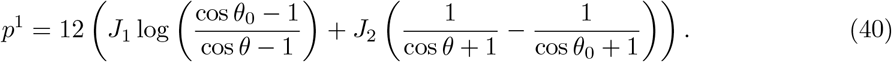

The velocity 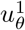 can be obtained from (36), while 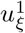 can be obtained from the continuity equation. Explicit expressions for *J*_*i*_ and *H*_*i*_, *i* = 1, 2 are given in §B.3

### 2.7 Numerical solution

In the case of a realistic brain with non-uniform displacements, we solve the system numerically using COMSOL Multiphysics^®^ version 6.4. Specifically, we first solve the leading order problem (21) on a rectangle (*θ, ϕ*) ∈ [0, *θ*_0_] × [0, 2*π*]. The mesh was generated by the software; we set the maximum element size to 0.05. We then solve the steady streaming problem (29) on the same domain, using the leading order solution to calculate functions in (30a).

We checked that the numerical solution agrees with the analytical solution for the spherical case with uniform displacements. We also performed a mesh test. In terms of maximum steady streaming velocity, the solution changed by 0.82 % after refining the mesh to have a maximum element size of 0.025 and by 0.94 % for 0.0125.

## 3 Pial surface geometry and displacements

### 3.1 Geometry

We used the geometry and displacement data from Adams *et al*. (2020) for 8 healthy subjects. For each subject, we were provided masks for grey matter (GM), white matter (WM) and CSF, which are probability maps obtained using SPM12 (Wellcome Centre for Human Neuroimaging, University College London; Adams et al., 2020). We start by reconstructing the brain geometry for each subject to obtain the function 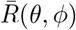. First, we apply a GM mask. We then manually place the centre 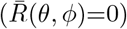 over the brain stem so that the outlet is centred on the same axis as *θ* = 0 and can be described with *θ* = *θ*_0_. Then, we interpolate the radial component of the surface on *θ*-*ϕ* grid of 200 ×400 points, using `scatteredInterpolant` function in Matlab R2024a (MathWorks) with the linear nearest neighbour extrapolation method. We then smooth the resulting surface using `imgaussfilt` function. This surface is then approximated with *N*_*g*_ spherical harmonics (*N*_*g*_ = 10). At this step, details of the brain sulci are neglected, and a ‘smoothed’ brain surface is obtained as an analytical function 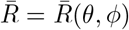 expressed in spherical harmonics. An example of a reconstructed brain is shown in the first panel of figure 3(a).

**Figure 3:**
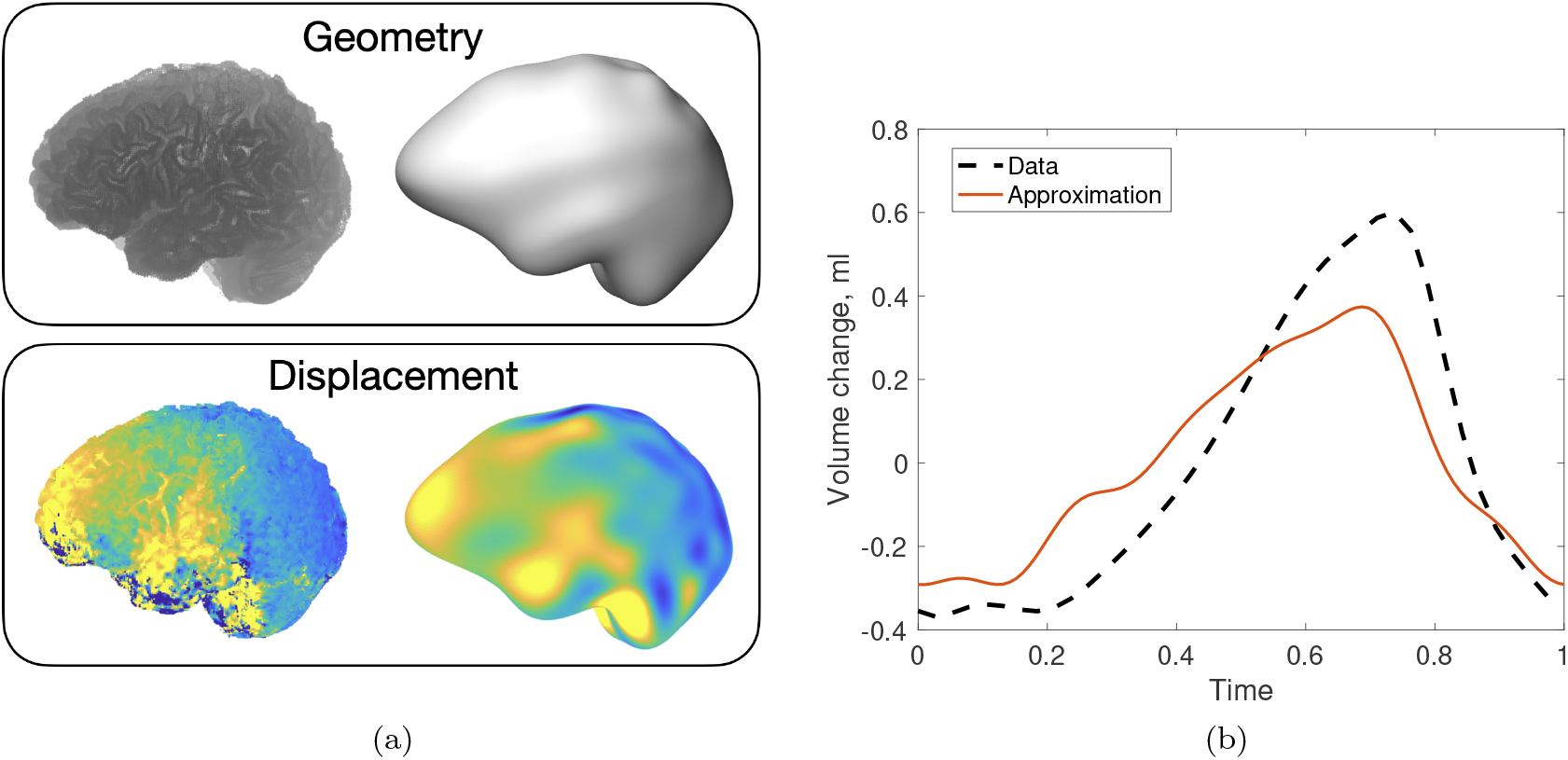
(a) Approximation of the geometry and surface displacement for Br19. Top panel: on the left, we show the cloud of points obtained after applying the GM mask to the data. On the right we show constructed analytical brain geometry with *N*_*g*_ = 10 spherical harmonics. On the bottom panel, we show brain displacements. The points on the left are coloured with the corresponding displacement in *z*-direction at time point 17. The figure on the right is the approximation described in §3.2. The colour scale is the same in both panels. (b) Resulting CSF volume change of brain Br19 measured in Adams *et al*. (2020) (dashed curve) and the model approximation during one cardiac cycle.

### 3.2 Displacement

To process the displacement data, we used the procedure and the codes developed in Karimi *et al*. (2023). The DENSE data covered 20 time points of one cardiac cycle for two opposite gradient polarities. We removed background errors by subtracting the data acquired with the negative polarity from those acquired with the positive polarity. The initial mask included GM and WM regions. That mask was eroded (three voxels) to remove regions near CSF, which have high uncertainty. The k-nearest neighbours algorithm (KNN, scikit-learn) (k = 150) was used to smooth the deformation data and reduce noise. To extract the deformation fields on the moving interfaces of interest, the masked data were extrapolated using nearest neighbour extrapolation and unstructured fast Fourier transform (FFT), developed by Karimi *et al*. (2023). In this work, we used 5 Fourier frequencies and the zero padding of 0.9.

After the smooth structured data was obtained, every time step of the data was approximated with *N*_*d*_ spherical harmonics. We selected *N*_*d*_ = 15, as choosing more harmonics led to overfitting and the appearance of non-physiological surface waves. We then approximated each coefficient of the expansion with the Fourier series with *N*_*h*_ = 6 harmonics in time. As a result, we obtained an analytical approximation of the displacement function in space and time, 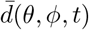. An example of surface displacement of one of the considered brains (named Br19, where the last two digits are from the subject’s pseudonym) in *z*-direction before and after the approximation is shown in figure 3(a).

In figure 3(b), we compare the volume change of brain Br19 computed from the approximation and measured in Adams *et al*. (2020). The model captures the qualitative behaviour of the volume change, but does not reproduce precise quantitative agreement. Using larger numbers of harmonics for 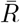 or for 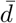 in space and time, or different smoothing in FFT, did not improve the agreement. We note, however, that our description does not capture changes in ventricular volume and the shape of brain sulci (which will locally increase the surface area), which are present in the measured volume change. We leave these for future investigation.

### 3.3 Spherical brain and uniform displacements

In this paper, we consider a simplified case of a spherical brain with uniform displacements, along with two other cases which combine simplified and realistic descriptions of geometry and displacements (described in table 3 and § 4.3). Below, we describe how we adapt geometry and displacement to such cases.

1. *Spherical brain*. We solve the system in spherical geometry. For this, we match the pial surface area in realistic and spherical brains by using the surface area of the brain 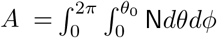for Br19 to compute the radius 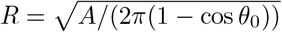 of the spherical brain. The dimensional values for these quantities are *Ā* = 0.07 m^2^ and 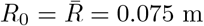. We note that this value is comparable to the radius of 0.066 m obtained from assuming spherical brain of volume 1200 ml (Cosgrove *et al*., 2007).
2. *Uniform displacement*. We also consider a case when the pial surface undergoes spatially uniform displacement. The surface velocity is then prescribed to reproduce the flux into the spine given by the blue dashed curve in figure 1(b) in §4.1.1 and §4.2.1, or a computed flux into the spine from Br19, reported for example in figure 8. We approximate these curves with 6 Fourier harmonics to obtain the expression 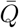. The velocity is then found from

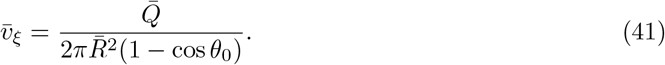

Consequently, 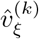 are the Fourier coefficients of the expansion of 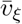 into 6 harmonics.

## 4 Results

### 4.1 Oscillatory flow

#### 4.1.1 Spherical case

In the case of a spherical brain with uniform displacements, we use the analytical solution computed in §2.6 for the surface velocity (41) constructed to obtain flux into the spine given by the mean curve in figure 1(b). The flow and pressure profiles in different time instances are shown in figure 4 and these times are annotated in figure 5(a). At the point of largest CSF flux, *t*_1_ (systolic phase), the flow goes into the spine (figure 4a). The velocity magnitude is always the largest at the outlet as it corresponds to the point of maximum flux (factor (1 −cos *θ*) in (33a)) and small surface area (factor 1*/* sin *θ* in (33a)). In figure 4(d), we show the corresponding pressure and the velocity at the outlet. The velocity has a flattened parabolic-like profile. The point of the highest pressure is at *θ* = 0, corresponding to the top of the head.

**Figure 4:**
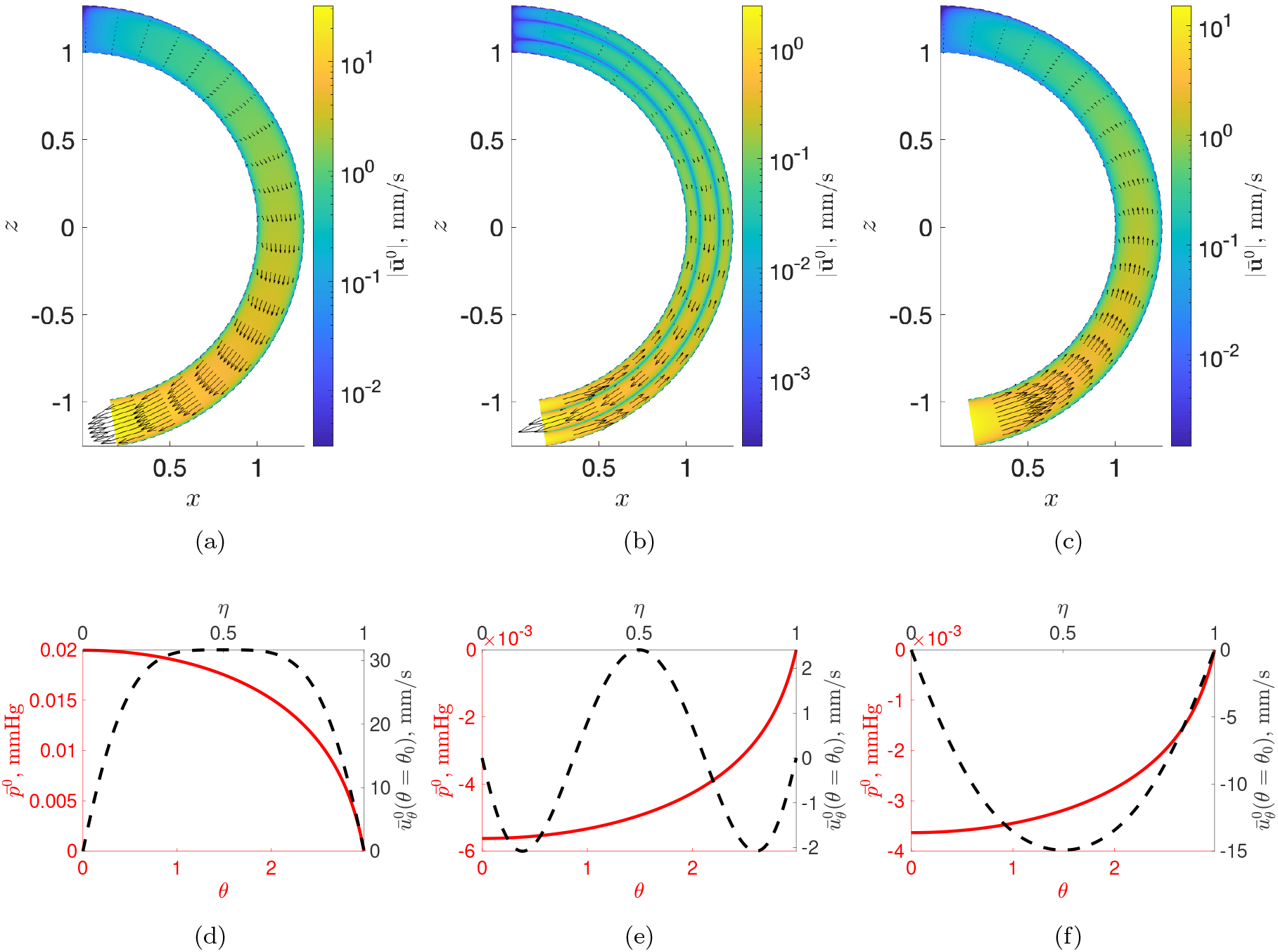
Flow profiles, pressures and outlet velocities in three different time steps. The first row is velocity fields (arrows) and velocity magnitude (colours) in the cross-section of the cSAS. The domain is axisymmetric with respect to *θ* = 0. The *η* component of the cSAS is magnified 10 times for better visualisation. Different columns represent different time steps: (a),(d) *t*_1_ = 0.88*T* (systolic phase); (b),(e) *t*_2_ = 0.11*T* (flow reversal); and (c),(f) *t*_3_ = 0.5*T* (diastolic phase), where *T* is the dimensionless period (*T* = 2*π*). The second row left axis: pressure (red solid lines) along the zenithal coordinate *θ* (bottom axis). Right axis: *θ* velocity 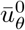 (black dashed lines) at the exit of the domain *θ* = *θ*_0_ as a function of *η* (top axis).

**Figure 5:**
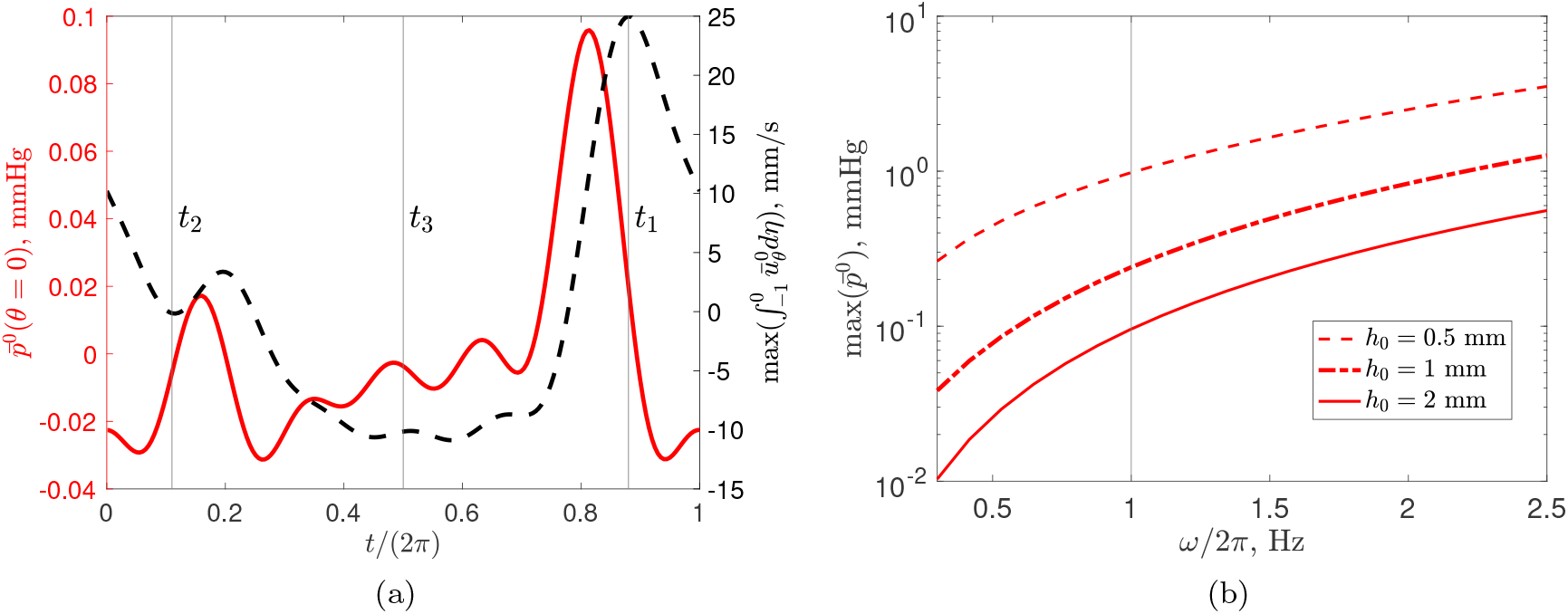
(a) Pressure at the top of the head *θ* = 0 (left axis) and depth-averaged velocity at the outlet (right axis) in SAS during the cardiac cycle. The vertical black lines represent time instances from figure 4. (b) Maximum pressure in cSAS as a function of the frequency of the oscillations for different thicknesses of cSAS. The vertical line corresponds to the reference heartbeat of 1 Hz.

The second time point *t*_2_ (figures 4b,e), is near flow reversal, thus, the net fluid flux is close to 0. The velocity profile changes direction with *η*, as the flow is directed into the cranium (in negative *θ* direction) near the pial and dura boundaries and into the spine in the middle.

At the third time point *t*_3_ representing diastolic relaxation, the flow is directed into the cranium. As a result, the pressure is negative at *θ* = 0.

Using eq. (33a), the depth-averaged velocity in the domain can be written as

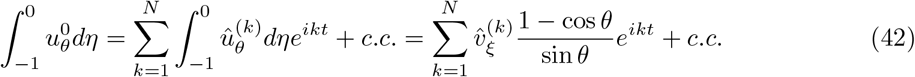

Its maximum value is shown in figure 5(a), right axis. The peak of the velocity coincides with the peak of the flow. According to eq. (42), the depth-averaged velocity does not depend on *α*. Therefore, owing to its scaling *h*_0_*ω*, the dimensional average velocity has a linear dependency on *ω* (not shown). The velocity and the pressure are also linear with respect to the velocity of the brain surface (see (33)).

The maximum pressure in the domain is reached at *θ* = 0. Its value during the cardiac cycle is shown in figure 5(a), left axis. The pressure peaks just before the velocity. The peak of the pressure as a function of the frequency of the heartbeat *ω/*2*π* is shown in figure 5(b) for different thicknesses of SAS *h*_0_. Unlike the case of small *α* (standard lubrication theory), pressure is not proportional to 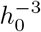.

#### 4.1.2 Realistic brain displacement

We now consider the realistic brain geometry and displacement described in §3.1 and §3.2. We first present the results for a single subject, Br19, and then study and compare eight subjects from Adams *et al*. (2020).

In figure 6 we show the flow profiles at different important time points, corresponding to systolic peak (first column, panels a,d,g,j), flow reversal (b,e,h,k) and diastolic relaxation (c,f,i,l). The same times points are annotated in figure 8(a), where we also show volumetric flow rate into the spine.

**Figure 6:**
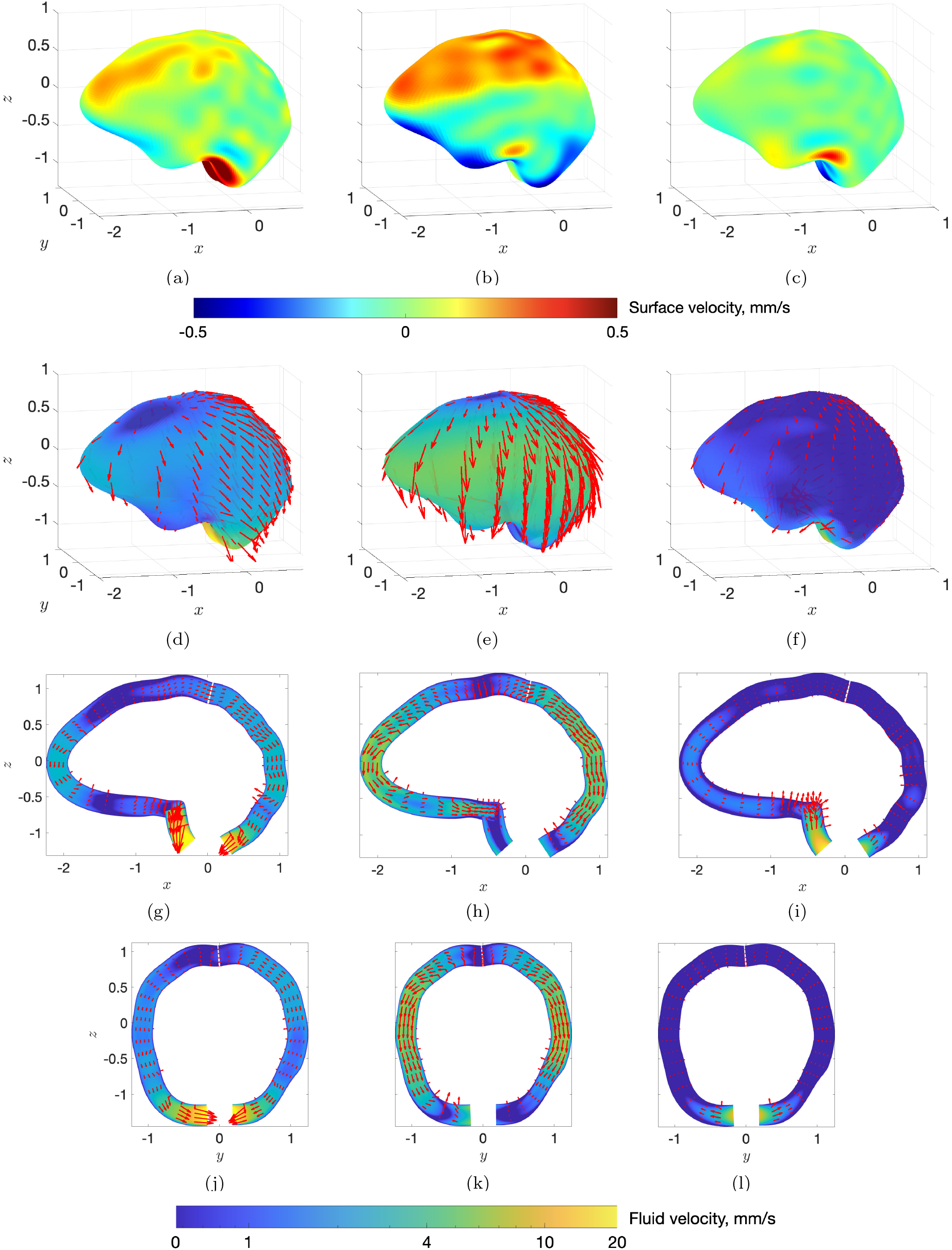
Flow profiles in three time instances (columns): *t*_1_ = 0.775*T, t*_2_ = 0.14*T, t*_3_ = 0.5*T*. The top row is the velocity 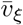 prescribed at the brain surface. The second row is the depth-averaged fluid velocity (arrows: velocity vectors; colors: velocity magnitude), while the third and fourth rows are fluid velocities in sagittal (*y* = 0) and coronal (*x* = 0) sections, respectively. In the coronal and sagittal sections, the domain appears to have non-uniform thickness, an artefact arising from the normal vector being misaligned with the plane. Furthermore, in the sagittal section, the non-uniqueness of spatial position in the region of higher curvature is an artefact of scaling used for visualisation (factor of 10 for *η*-component).

During the systolic phase (*t*_1_ = 0.775*T*, first column), the flow is directed towards the spine. The depth-average profile (d) shows the effect of the pial surface velocity (a): the flow is directed towards the regions of the expansion of the domain (negative 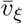). The outward movement of the brain surface (positive 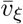) drives the fluid flow, with the region of zero flow closer to the frontal part of the brain (instead of *θ* = 0 as in the sphere). The velocity is the largest at the outlet *θ* = *θ*_0_, as seen in sagittal and coronal cross-sections (g,j).

During the flow reversal stage (*t*_2_ = 0.14*T* second column) the top part of the pial surface is moving outwards and the bottom part moves inwards (b). This results in almost no net flow into the spine. However, the velocity magnitude in the domain is still quite large, about 5 mm/s (e,h). Finally, during diastolic relaxation (*t*_3_ = 0.5*T*, second column) the pial surface exhibits high spatial variability with movement both inwards and outwards (c). However, the overall flow direction is from the spinal SAS to cSAS (f,i,l). The video of surface velocity and the resulting depth-integrated fluid velocity during one cardiac cycle can is shown in Supplementary Movie 1.

In figure 7(a-c) we show the pressure distribution in the domain in the same time points as in figure 6. We recall that we prescribe zero pressure at the boundary with the spine. In *t* = *t*_1_, pressure at the top of the head is around 0.1 mmHg. In *t* = *t*_2_, the pressure is negative around the cribriform plate and close to zero at the top of the head. In *t* = *t*_3_, pressure variations are quite small, which is consistent with almost no flow in the domain (fig. 6f).

**Figure 7:**
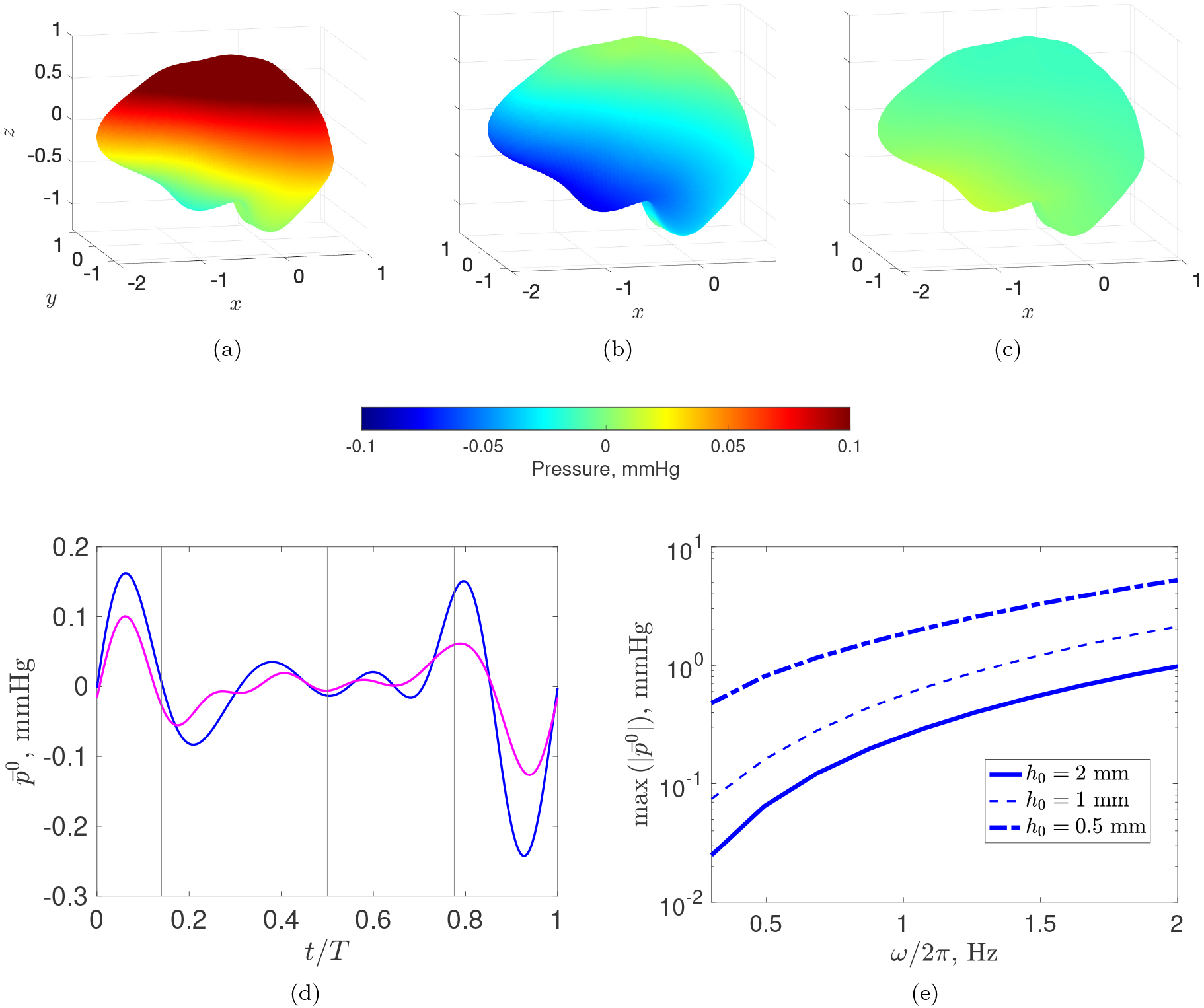
(a-c) Pressure distribution in the domain at the same time points as Fig. 6. (d) Pressure in *θ* = 0 (blue) and mean pressure in *θ* = *π/*2 (magenta), calculated as 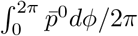, as functions of time. (e) Maximum pressure as a function of *ω*. Different curves represent different cSAS thickness.

In figure 7(d), we show the pressure near the top of the head *θ* = 0 and the average pressure in the equatorial plane (*θ* = *π/*2) as functions of time. At *θ* = 0, the pressure peaks after the flow peak, and there is a large pressure change from the systolic phase to flow reversal, from -0.24 mmHg to 0.16 mmHg. The maximum absolute value of pressure in the cSAS over time is about 0.26 mmHg, located around the top of the head.

Figure 7(e) shows maximum pressure in the domain as a function of *ω*. As in figure 5, the pressure increases with *ω*, reaching values of about 3 mmHg for thin cSAS and large frequencies.

The outflow into the spine is shown in figure 8(a). With a dashed line, we show the PC MRI measurements on the same subject. Our model gives a reasonable qualitative agreement with the measurements. However, it underpredicts the magnitude of the systolic phase, and the flow reversal region (between *t* = 0.05*T* and *t* = 0.2*T*) is not well captured.

**Figure 8:**
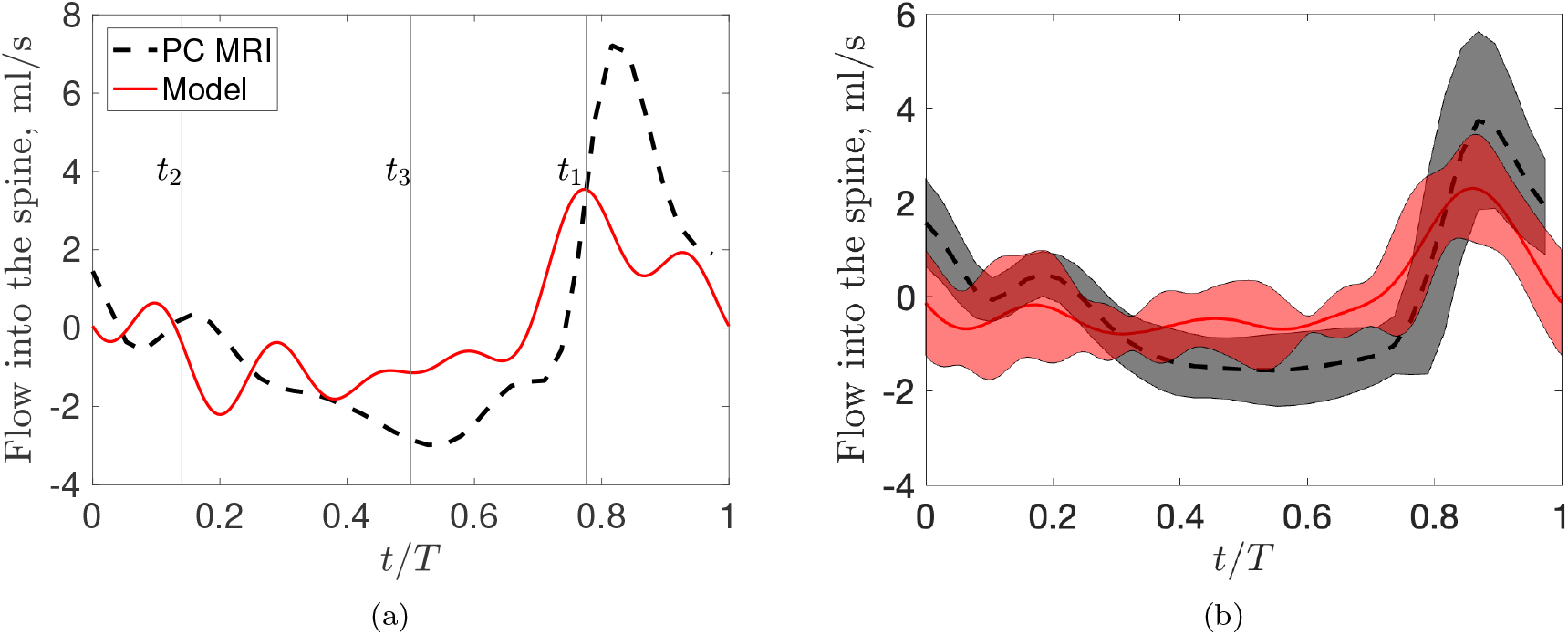
CSF flow into the spine computed (red) as 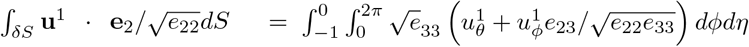. The black dashed line shows PC MRI measurement of flow into the spine for the same subject (Adams *et al*., 2020). (a) From Br19 and (b) averaged between 8 subjects. In (b), the shading around the mean curve represents standard deviation. In (a), vertical lines show times *t*_1_, *t*_2_ and *t*_3_ considered in figures 6 and 7.

#### 4.1.3 Comparison between subjects

To assess the difference between the subjects, we first show the maximum spatial pressure drop in the domain for 8 subjects from Adams *et al*. (2020) in the second row of table 2. We note that this is not the temporal intracranial pressure variation known as the ‘pulse pressure’, but rather the spatial pressure drop along the cSAS. The mean value is 0.23 ±0.1 mmHg. We also report the maximum oscillatory velocity at the equatorial plane (third row). There is some variability between subjects, with a mean value of 9.75 ± 4.3 mm/s.

**Table 2:** Comparison for maximum pressure drop in the domain and steady streaming velocity between different brains. The first row is the subject number. The second row is the maximum pressure drop (in space and time) in the domain. The fourth row is the maximum depth-averaged leading order velocity at the equatorial plane (*θ* = *π/*2). The fourth and fifth rows denote maximum depth-averaged steady streaming velocity at equatorial plane *θ* = *π/*2 and at the top of the head *θ* = 0, respectively. In bold, we denote the Br19 considered in the result section. The last column shows the mean and the standard deviation between the brains.

| Subject number | Br18 | **Br19** | Br20 | Br21 | Br22 | Br34 | Br35 | Br46 | mean |
| --- | --- | --- | --- | --- | --- | --- | --- | --- | --- |
| $\max \vert \bar{p}^0\vert $, mmHg | 0.42 | 0.26 | 0.15 | 0.2 | 0.34 | 0.11 | 0.13 | 0.23 | $0.23 \pm 0.1$ |
| $\max \left\vert \int_{-1}^0 \bar{\mathbf{u}}^0\vert _{\theta=\frac{\pi}{2}} d\eta \right\vert $, mm/s | 16.2 | 12.2 | 8.2 | 7.3 | 14.3 | 3.3 | 6.5 | 10 | $9.75 \pm 4.3$ |
| $\max \left\vert \int_{-1}^0 \langle \bar{\mathbf{u}}^1 \rangle\vert _{\theta=\frac{\pi}{2}} d\eta \right\vert $, $\mu\text{m/s}$ | 39.6 | 35.8 | 16 | 40.5 | 13.6 | 3.72 | 16.1 | 27.2 | $24.1 \pm 13.7$ |
| $\left\vert \int_{-1}^0 \langle \bar{\mathbf{u}}^1 \rangle\vert _{\theta=0} d\eta \right\vert $, $\mu\text{m/s}$ | 25.9 | 18.5 | 8.1 | 18.5 | 2.38 | 9.02 | 19.2 | 14.5 | $14.5 \pm 7.6$ |

The mean outflow for 8 brains is shown in figure 8(b). Again, there is a qualitative agreement with PC MRI. Possible reasons for quantitative disagreement are discussed in §5.

### 4.2 Steady streaming

#### 4.2.1 Spherical case

The oscillatory flow reported in the previous section induces steady streaming at 𝒪 (*ϵ*) (§2.6.2) shown in figure 9(a). The flow consists of three regions across the thickness of the domain: near the boundaries it is directed towards the spine, while at the middle of the domain, it is directed towards *θ* = 0. The velocity profile at the outlet *θ* = *θ*_0_ is shown with a black dashed line in figure 9(b) (right axis).

**Figure 9:**
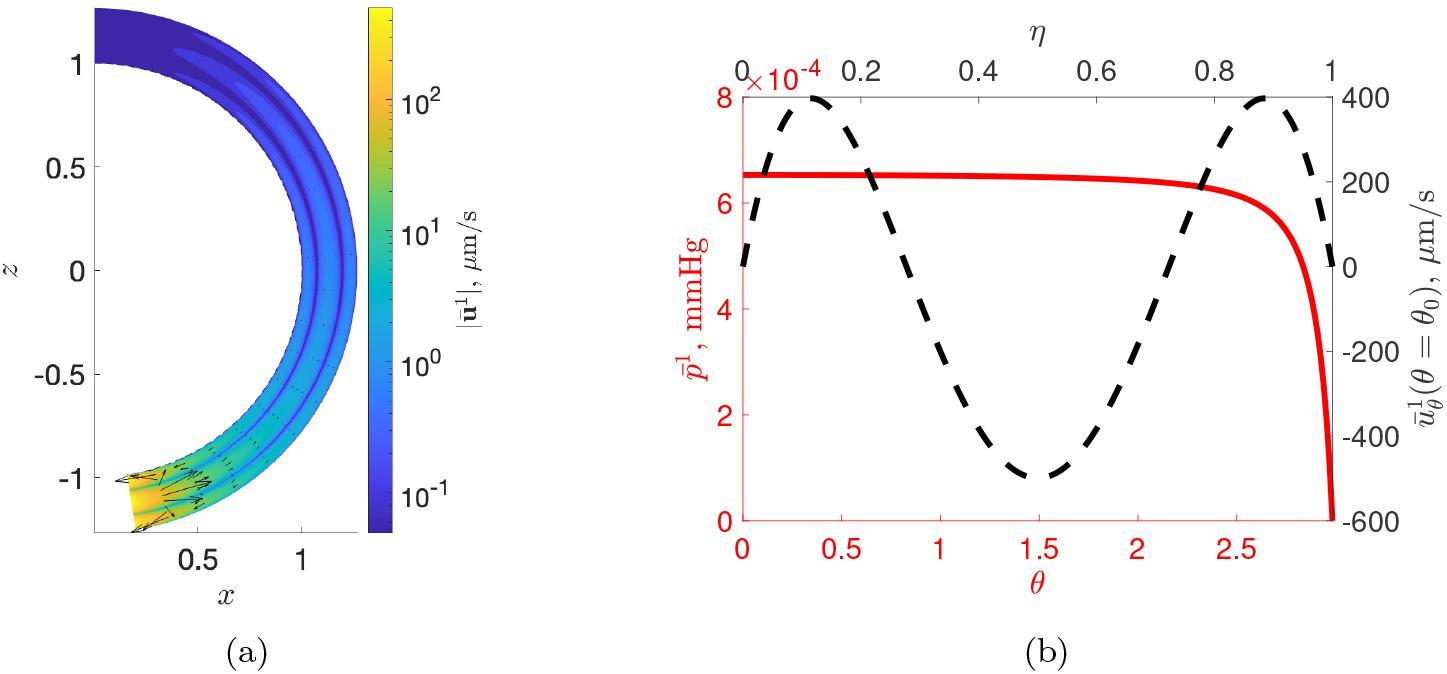
(a) Steady streaming velocity field (arrows) and velocity magnitude (colours) in the cross-section of the cSAS. The *η* component of the cSAS and 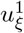 velocity are magnified 10 times for better visualisation. (b) Left axis: steady streaming pressure 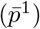 along the zenithal coordinate. Right axis: *θ*-velocity component, 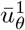 at the exit of the domain *θ* = *θ*_0_.

The corresponding pressure along the domain, given by (40), is shown with a red line in figure 9(b), left axis. The pressure remains almost constant in the domain and drops to 0 at the outlet.

The velocity magnitude at the outlet is about 0.5 mm/s. This is at least an order of magnitude smaller than the peak velocity of oscillatory flow reported in §4.1.1 (≈ 1 cm/s). The rest of the domain, however, has much smaller steady streaming velocities, with a mean value in *η* = − 1*/*2 being 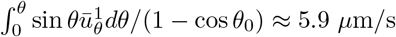.

#### 4.2.2 Realistic brain displacement

Steady streaming flow for the realistic case is shown in figure 10(a). The velocity is largest at the outlet with a maximum value of 6.3 mm/s. Interestingly, there is a net circulation flow from the front of the head to the back. The maximum depth-averaged velocity magnitude at *θ* = 0 is 18.5 *μ*m/s and at *θ* = *π/*2 is 35.8 *μ*m/s.

**Figure 10:**
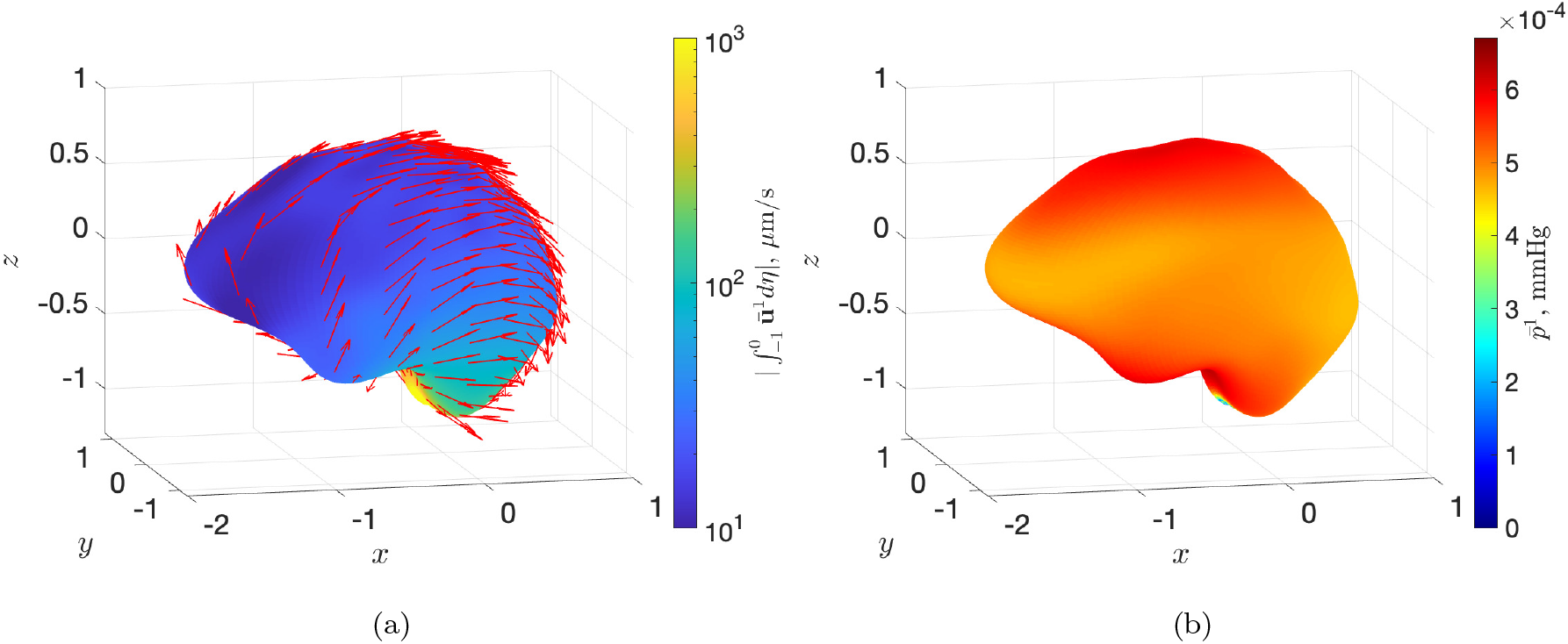
(a) Depth-averaged steady streaming normalised velocity vectors (arrows) and magnitude (colours) in the cross-section of the cSAS. (b) Steady streaming pressure 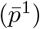 in the domain. Both figures are from Br19.

The corresponding steady streaming pressure is shown in figure 10(b). A large pressure gradient is found around the outlet, similar to the spherical case in figure 9(b).

#### 4.2.3 Comparison between subjects

In our model, all 8 subjects exhibit front-to-back steady circulation, similar to the one in Br19 (results not shown). Maximum steady streaming depth-averaged velocity is reported in table 2 in *θ* = 0 and *θ* = *π/*2. There is some variability between subjects, with the mean value of 24.1 ±13.7 *μ*m/s at *θ* = *π/*2.

### 4.3 The role of brain geometry and displacement

We now study the effect of brain geometry and brain displacement for Br19. We construct four case studies outlined in table 3, which consider combinations of spherical and realistic (from §3.1) geometry with uniform and realistic (from §3.2) displacement. We chose the radius of the spherical brain, *R*_0_, to have the same surface area as in Br19 (see §3.3). To have the same volume change in all cases, the uniform displacement in cases 1 and 3 is chosen to reproduce the flow rate from the red line in figure 8(a). For case 2, the surface velocity is multiplied by N*/* sin *θ*, to account for differences in local areas in different geometries. This way, all the cases have the same flux into the spine.

**Table 3:** Different case studies considered in figures 11 and 12. *R*_0_ denotes constant radius, representing spherical geometry and *d*(*t*) represents spatially uniform displacements.

|  |  |  |  |  |
| --- | --- | --- | --- | --- |
|  | Case 1 | Case 2 | Case 3 | Case 4 |
| Brain radius | $R_0$ | $R_0$ | $R(\theta, \phi)$ | $R(\theta, \phi)$ |
| Surface displacement | $d(t)$ | $d(\theta, \phi, t)\mathbf{N}/\sin\theta$ | $d(t)$ | $d(\theta, \phi, t)$ |

#### 4.3.1 Leading order pressure

The pressure at *θ* = 0 during the cardiac cycle for each case is shown in figure 11(a). The pressures for uniform displacement in cases 1 and 2 are almost identical. The maximum values in the overall domain are 0.106 and 0.108 mmHg for cases 1 and 3, respectively. This suggests that in the case of uniform displacement, the specific brain geometry does not play a significant role. The realistic displacements of the domain change the pressure profile, as can be seen from the figure. The pressures in cases 2 and 4 almost overlap, with the maximum pressure magnitude of about 0.25 mmHg. This indicates that the maximum pressure drop is governed by displacement, not geometry (as long as the surface areas are the same). Therefore, to study leading-order pressure, the brain can be approximated as a sphere with non-uniform displacements.

**Figure 11:**
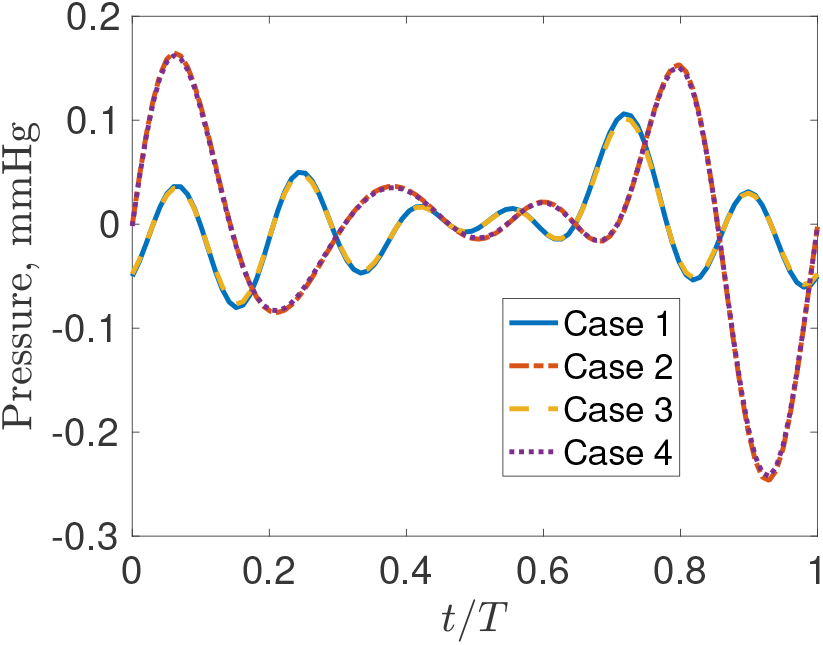
Pressure at *θ* = 0 during cardiac cycle for different cases from Table 3.

#### 4.3.2 Steady streaming

It is of interest to see whether the transversal steady streaming is an effect of the geometry or of the spatially heterogeneous surface velocity. In figure 12, we show steady streaming in cases 2 and 3 from table 3 (with case 1 shown in fig. 9(a) and case 4 in fig. 10(a)). In both cases, the circulation around the brain is present, with magnitudes being larger in case 2. This suggests that both geometry and non-uniform displacements play a role in the formation of net flows in the steady streaming.

**Figure 12:**
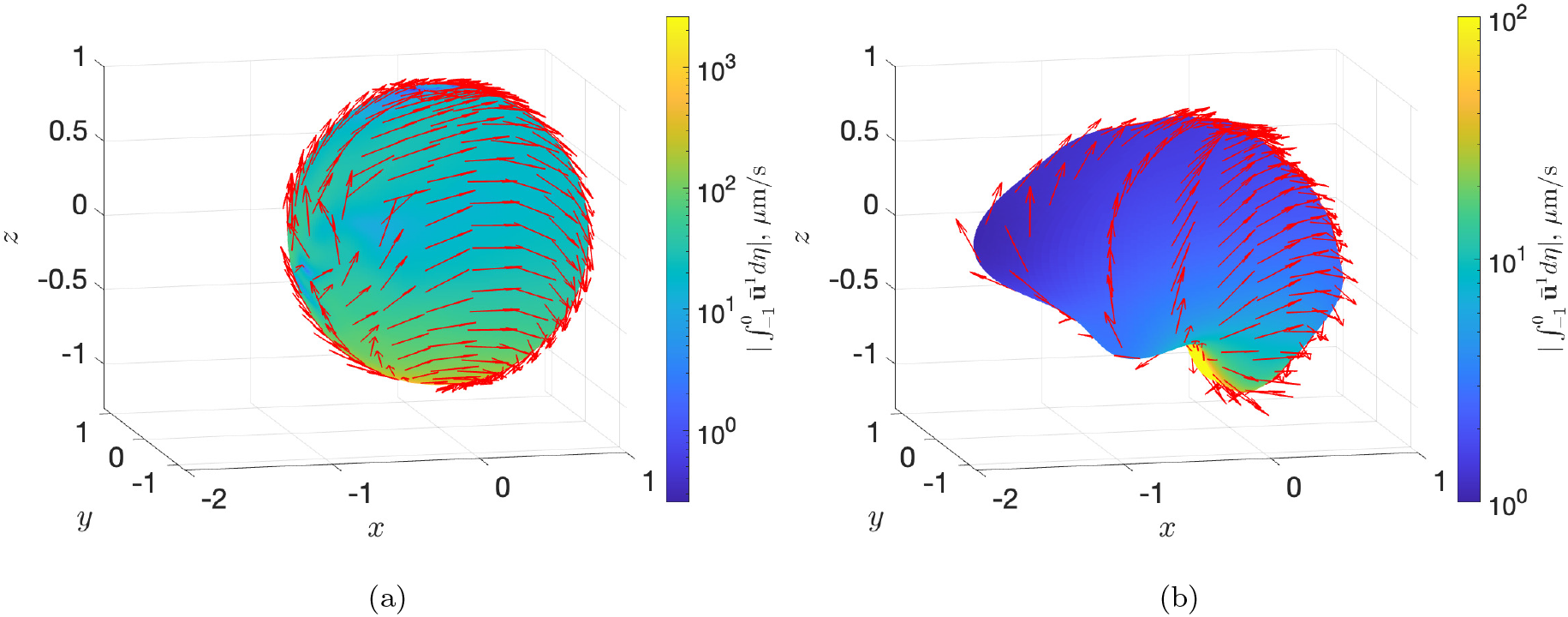
Steady streaming for the cases (a) 2 and (b) 3 from the table 3.

## 5 Discussion

In this work, we developed a reduced model for cerebrospinal fluid (CSF) flow in cranial subarachnoid space (cSAS) driven by brain surface displacements during the cardiac cycle. Taking advantage of the small aspect ratio of the domain *ϵ*, we used lubrication theory to solve the problem both for the ideal case of a spherical brain with uniform displacement and for the realistic case, where we approximated brain geometry and displacements from DENSE MRI data with spherical harmonics. Our reduced model allows the exploration of flow mechanisms and the role of model parameters.

The leading-order flow is purely oscillatory and in phase with the brain’s surface velocity. It generates an oscillatory pressure gradient within the cSAS. Our model predicts a maximum spatial pressure drop along the cSAS of 0.23± 0.1 mmHg (averaged across 8 subjects). This is in good agreement with the pressure drop in cSAS of about 0.21 mmHg computed with the fluid-structure interaction model of Causemann *et al*. (2022). Larger pressure drop is found for thinner cSAS and for higher heartbeat frequency. This spatial variation is much smaller than the intracranial pressure (ICP) itself (about 5–15 mmHg in healthy adults (Yuh & Dillon, 2010)) and smaller than its cardiac-cycle variation (pulse pressure), which is approximately 4 mmHg (Eide, 2021; Causemann *et al*., 2022).

Our model predicts the maximum velocity of the oscillatory flow in the equatorial plane of 9.95±4.3 mm/s. This is comparable with CSF velocities in numerical studies of Gupta *et al*. (2010), where the authors report a value of 12.6 mm/s, and Causemann *et al*. (2022), where the peak velocity of about 10 mm/s in cSAS was reported. These values, however, are much larger than the velocities measured in van der Voort *et al*. (2025). Indeed, using the measured CSF displacements of about 25 *μ*m (Figure 10 in their manuscript) over ∼0.25 s of systolic phase leads to velocities of order 100 *μ*m/s. We anticipate that this discrepancy may be due to the absence of sulci (brain folds) in our model and the assumptions we made on cSAS thickness, however, further investigation is required.

At order *ϵ* we find that there is a steady streaming flow. For a spherical brain with a uniform surface displacement, this steady streaming produces zero net flow: the velocity is directed towards the spine near the boundaries of the domain (pial surface and dura membrane), and towards the top of the head in the middle region. This flow is similar to those found in classic works on steady streaming in blood vessels (Secomb, 1978; Hydon & Pedley, 1993) and, more recently, in the spine with axisymmetric geometry (Sánchez *et al*., 2018).

In the case of a realistic brain with displacements from DENSE MRI, our model predicts that there is a steady front-to-back circulation for all 8 subjects: the fluid travels upwards (towards the top of the head) in the anterior part of the cSAS and downwards in the posterior part. To our knowledge, this is the first report of such circulation in cSAS. Recent analytical model of the CSF flow in the spine, approximated as the space between two eccentric cylinders (Sánchez *et al*., 2018), found a similar circulation there, though in the opposite direction: the fluid travels downwards in a thicker space in the anterior spinal SAS and upwards in a thinner posterior space. Coenen *et al*. (2019) then used patient-specific geometry to find Lagrangian mean velocity (which includes steady streaming and Stokes drift). They show that the direction of the circulation depends on the position of the spinal cord in the spinal canal, and it varies between subjects. Future investigation is needed to understand how (and if) spinal and cranial circulations co-exist and interact with each other.

The causes of the circulations are also different: the circulation in the spinal SAS is due to the spatial heterogeneity of the domain, while that in cSAS is governed by the shape of the brain and its non-uniform pulsations. We note that there is evidence that cSAS is spatially non-uniform, with thicker space in the front and thinner in the back (Frydrychowski *et al*., 2012). However, owing to MRI resolution and noise, it was challenging to reconstruct cSAS geometry from our data, and we kept the uniform thickness of the cSAS in this work, leaving the effect of spatial heterogeneity of the cSAS for future investigation.

The steady streaming flow is about 24.1 ± 13.7 *μ*m/s (averaged across 8 subjects), which is similar to the flow expected from CSF production in the choroid plexus and drainage to the superior sagittal sinus (4.2 *μ*m/s van der Voort *et al*. (2025)). However, novel MRI measurements of net CSF flow by van der Voort *et al*. (2025) do not detect any net flow of expected magnitude towards the superior sagittal sinus. This means that our front-to-back steady streaming circulation is also not detected. While more imaging studies are needed to investigate the presence of this circulation, we note that we may also be overestimating the steady streaming velocities, since we treat cSAS as an open space, though in reality, it is a fibrous space containing arachnoid trabeculae (Saboori & Sadegh, 2015). Indeed, Nozaleda *et al*. (2025) found that steady streaming generated by oscillatory pressure gradient is smaller in a porous channel than in an open channel. The effect of fibers could be accounted for by including porous effect modeled for instance by Brinkmann’s equation (Gupta *et al*., 2009, 2010), which will be the subject of future research.

Steady streaming found in this work may contribute to solute transport, including clearance of metabolic waste from the brain and drug delivery. Indeed, for large solutes like amyloid-beta with diffusion coefficient *D* = 1.8 × 10^−10^ m^2^/s (Waters, 2010) and steady streaming velocity of *U*_*ss*_ = 10 *μ*m/s, the corresponding Péclet number is *Pe* = *U*_*ss*_*h*_0_*/D* ≈110. In the spherical case with uniform displacements, the mean speed in the midline of the domain is about 5.9 *μ*m/s. Although there is no net flow, the circulations in different directions across the cSAS thickness will enhance solute transport. In the presence of front-to-back circulation, the net flow will drive solute transport in the flow direction. If CSF turnover flow is present (Smets *et al*., 2024), it will act in concert with the steady streaming to transport solutes. In the absence of drainage into the superior sagittal sinus, the steady circulation predicted by our model could facilitate the transport of CSF to alternative exit routes.

The oscillatory flow may also contribute to net solute transport via Stokes drift, when spatial variations in oscillatory velocity cause fluid particle displacements to accumulate into a net drift over successive cycles (Larrieu *et al*., 2009). Lawrence *et al*. (2019) showed that in spinal SAS, Stokes drift is much smaller than steady streaming. With a toy model of cSAS consisting of a plane channel with uniform displacements, we also recently showed that there is an order of magnitude separation between Stokes drift and steady streaming (Neff *et al*., 2026). Nevertheless, the Stokes drift may become more significant when the brain surface is curved (e.g. in the presence of sulci) and for non-uniform displacements.

Our analytical model provides insight into CSF flows in cSAS. There are several limitations, however. First, the leading order flow does not quantitatively agree with the measured flow via PC MRI. We attribute this to the fact that we do not include sulci, which locally increase CSF displacement, and thus we underpredict the CSF flux. Additionally, there is scope to improve our data reconstruction and smoothing algorithms.

Second, our reconstruction does not include all anatomical details at the base of the brain and cerebellum, as lubrication theory requires smooth geometry. These effects may be added in future development of the model.

Third, we do not account for the possible deformation of the brain surface, modelled as rigid, in response to fluid pressure. To estimate whether we generate enough pressure for such deformation, let us assume a linear relationship between pressure 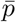 and height 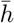 in cSAS, 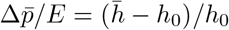, where *h*_0_ is the reference thickness. The elastic modulus of the brain at the frequency of the heartbeat is estimated to be around *E* ≈ 30 Pa (Herthum *et al*., 2021; Burman Ingeberg *et al*., 2025). To displace parenchyma by Δ*h* = *ϵ h*_0_ (comparable to the observed displacements), one needs to apply a pressure of Δ*p* = *ϵ E* ≈ 0.8 Pa ≈ 0.006 mmHg, which is smaller than the leading order pressure found in this work. Therefore, we may expect that the brain deforms as a result of CSF motion. However, since brain surface motion in our model is informed by *in-vivo* measurements, we expect that brain deformation by CSF flow is already present in the data and thus does not need to be included explicitly in the present model.

Finally, we neglected the exchange between the CSF and the interstitial fluid and the CSF in the perivascular spaces (PVS). The exchange with the interstitial fluid was recently estimated to be of the order of nanoliters per second, which is about 6 orders of magnitude smaller than the pulsating CSF flow (Causemann *et al*., 2022). This effect can then be safely neglected. However, the exchange with PVS is likely to be more significant, as shown in recent MRI tracer studies Eide & Ringstad (2024). This exchange is important due to its implications for the clearance of solutes, and will need to be addressed in future studies.

## Acknowledgments

The work of MD and AG was supported by the Engineering and Physical Sciences Research Council grant EP/R020205/1 to AG. JJMZ acknowledges the support of the Dutch Research Council (NWO, project number 18674). The authors would like to express the deepest gratitude to Fariba Karimi for providing the Python code to project MRI data onto a given brain surface. We also thank Dr Hadrien Oliveri, Andrew Ahern and Alannah Neff for very helpful discussions about the model.

## A Derivation of the equation in curvilinear coordinates

### Coordinate system

In this section, we derive the necessary expressions for a curvilinear coordinate system derived. We introduce a spherical-like surface **s**, parametrised with *θ* ∈ [0, *π*] and *ϕ* ∈ [0, 2*π*], as in spherical coordinates, so that **s** = [*R*(*θ, ϕ*) sin *θ* cos *ϕ, R*(*θ, ϕ*) sin *θ* sin *ϕ, R*(*θ, ϕ*) cos *θ*]. The function *R* is taken from MRI images and non-dimensionalised by *R*_0_, as explained in the main text. We then have two tangent vectors

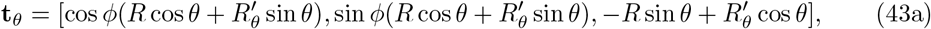

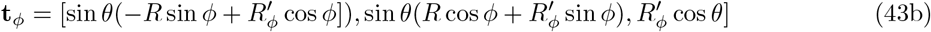

and the normal vector to the surface 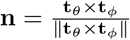. For a thin space around the surface and assuming that curvature is always much smaller than thickness, we can write the position vector **r** as

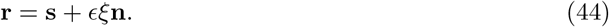

Neglecting order *ϵ* terms, we introduce a coordinate system (*x*_1_, *x*_2_, *x*_3_) = (*ξ, θ, ϕ*) in a thin region around the surface **s** and define base vectors as

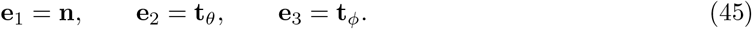

Note that this base is not orthonormal, as 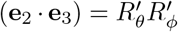. However, by construction, **e**_1_ is normal and orthogonal to **e**_2_ and **e**_3_. The metric tensor for this basis reads

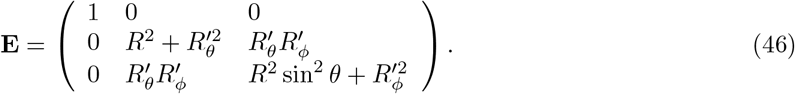

Each *i, j* component of this tensor is computed as *e_ij_* = (***e**_i_ · **e**_j_*). The velocity components can be written as 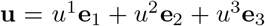, or alternatively in terms of unit vectors, 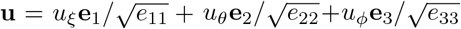. The transformation between new basis and unit vectors (**i, j, k**) in Cartesian system can be written as

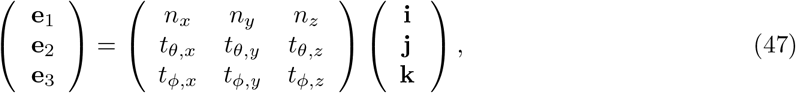

where subscripts *x, y, z* indicate corresponding components of vectors (45). Note that the matrix in (47) is the transpose of a Jacobian of the position vector (44), with the first row scaled by 1*/ϵ* and terms of order *ϵ* omitted in rows 2 and 3.

For convenience, we also introduce the conjugate or reciprocal basis

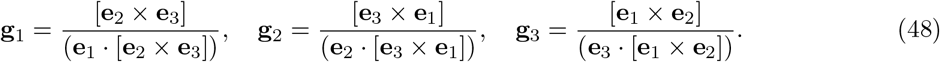

The conjugate metric tensor with components *g*_*ij*_ = (**g**_*i*_ · **g**_*j*_) will then read

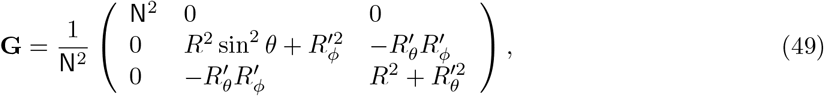

where

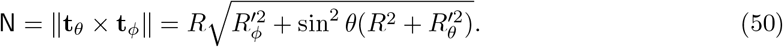

By construction, (**e**_*i*_ · **g**_*j*_) = *δ*_*ij*_, where *δ*_*ij*_ is Kronecker delta. The matrix formed from column vectors (**g**_1_, **g**_2_, **g**_3_) is inverse of the matrix in (47) and thus transforms (**e**_1_, **e**_2_, **e**_3_) back to (**i, j, k**). With this, the velocity vector can be written as

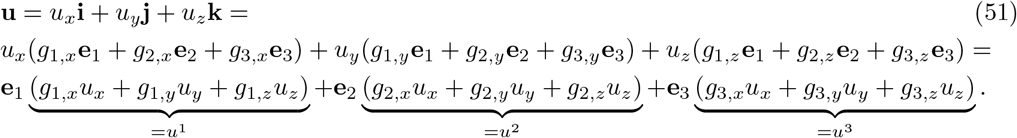

### The equations in curvilinear coordinates

The dimensional Navier-Stokes equations in non-orthogonal curvilinear coordinates read (Hill & Stokes, 2018)

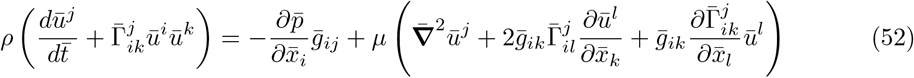

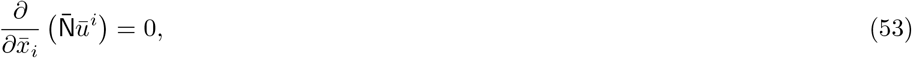

where we assume Einstein’s summation. The velocity components ū^*i*^ are related to physical velocity components ū_*ξ*_, ū_*θ*_, ū_*ϕ*_ as 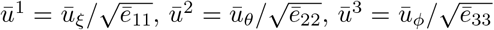, where ē_*ii*_ (no summation) are dimensionalised diagonal components of metric tensor (46). 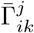 are dimensional Christophel symbols of the second kind, which are evaluated as 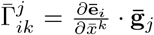. The material derivative 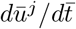 and the Laplacian can be written as

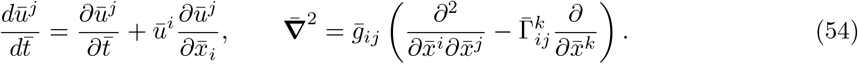

### Lubrication theory

We now will follow the standard lubrication theory simplification. Based on the construction of our coordinates, the *ξ* component will always be orthogonal to the surface and *ϵ* times smaller than the other components. Using the scaling from (2.3) we non-dimensionalise the equations (52). We obtain two dimensionless groups reported in (2.3). Following standard lubrication assumptions, we neglect the contributions of order *ϵ* ^2^ and *ϵ* ^2^*/α*^2^ and smaller and obtain equations (10). In that expression, the functions *ζ*_*i*_, *i* = 1, …, 4 are as follows

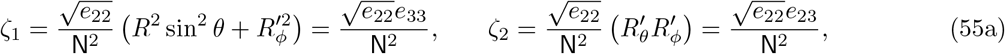

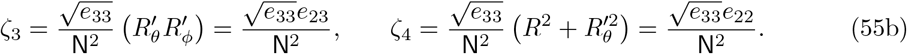

The inertial terms of order *ϵ*, 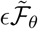 and 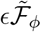, are reported in (11), and their non-linearity will generate steady streaming at the order *ϵ*. The residual terms in (10b), (10c), 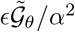 and 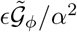, are also of the order *ϵ*. They originate from the viscous terms and the pressure terms. These terms are linear (at the order *ϵ*) with respect to velocity and pressure and will not contribute to the steady streaming at the order *ϵ* .

Note that for a spherical surface *R*(*θ, ϕ*) = 1, the pressure gradient coefficients are *ζ*_2_ = *ζ*_3_ = 0, *ζ*_1_ = 1 and *ζ*_4_ = 1*/* sin *θ*, which is consistent with lubrication equations in spherical coordinates (see e.g. Dvoriashyna *et al*. (2019)). Note that in that reference, there is *ϵ r* term multiplying the pressure gradient, which we do not have here. This is because we neglected *ϵ* terms in our base vectors (45). Since we are only interested in the leading-order solution and first-order steady streaming, these terms would not affect our system.

To calculate flux into the spine at *θ* = *θ*_0_, we use the following expression

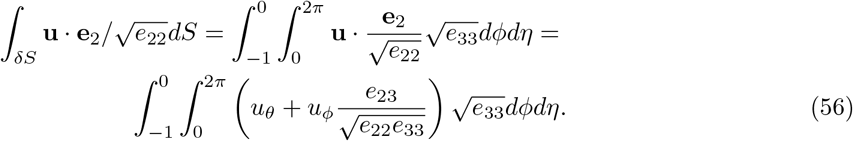

## B Model functions

### B.1 Inertial terms and functions for leading order solution

In this section we will report inertial terms, functions *f* ^(*k*)^, *F* ^(*k*)^, 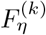. After the change of variables 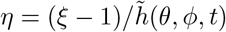, where 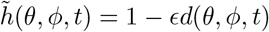, the inertial terms read

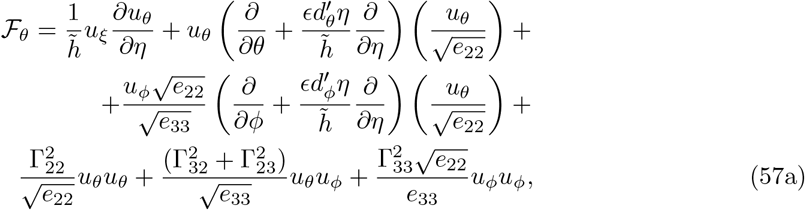

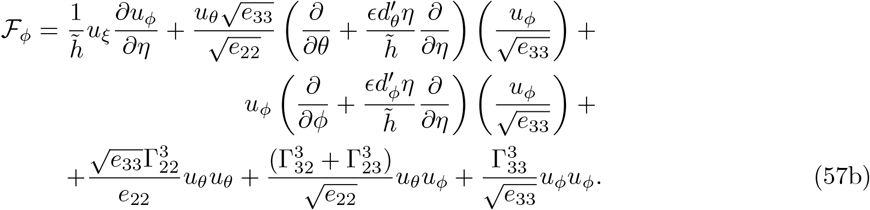

Here, 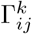 are Christoffel symbols, which we report in §B.4.

Recalling that 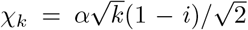, we can write the functions appearing in the velocity and pressure expressions (20),(21) and (23) as follows

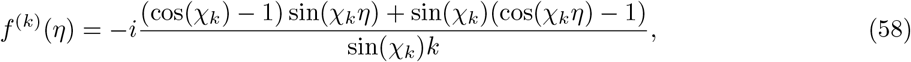

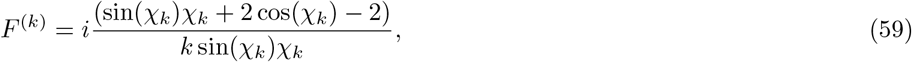

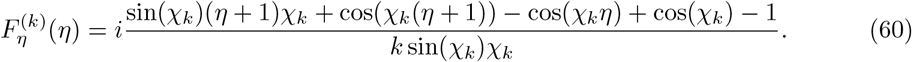

### B.2 Functions and constants in the steady streaming problem

For the convenience we will denote functions from (20) as 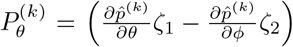 and 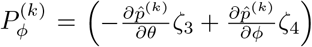. The function 𝒮_*θ*_ from (27), can be written as follows

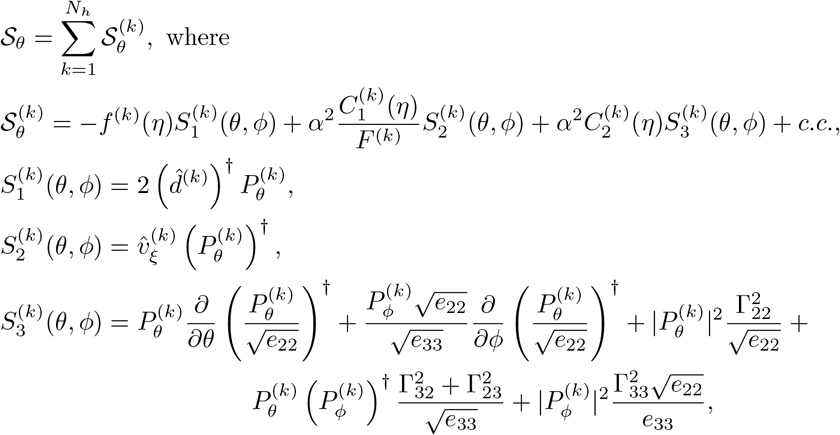

with 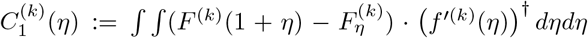, and 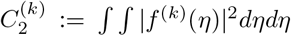. The expressions for these functions are reported below

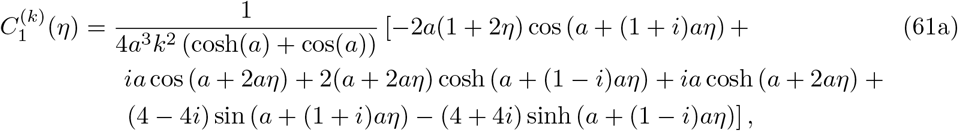

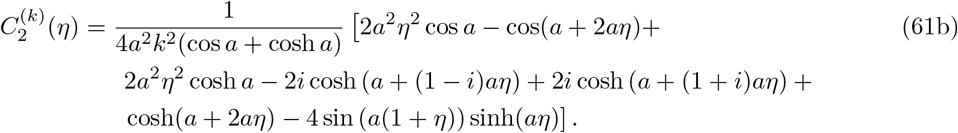

The functions 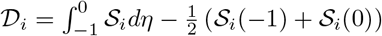 in (29) can be written as follows

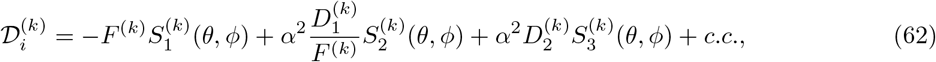

where

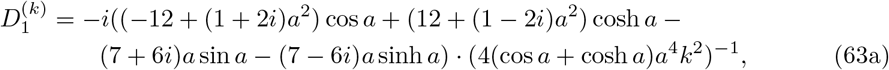

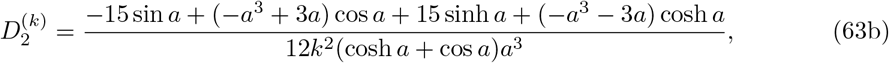

where 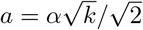.

The simplified expressions for 𝒦_*i*_, *i* = *θ, ϕ* in (29) are

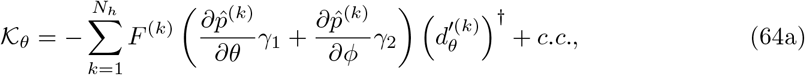

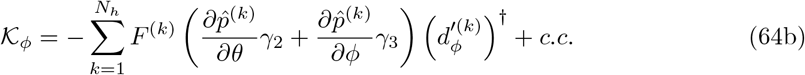

Furthermore, the integral in the first term in (29) is

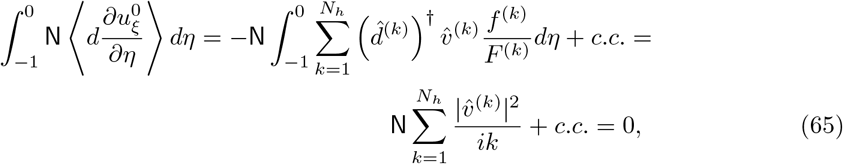

since the last expression has zero real part.

The normal velocity 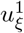is found from the continuity equation

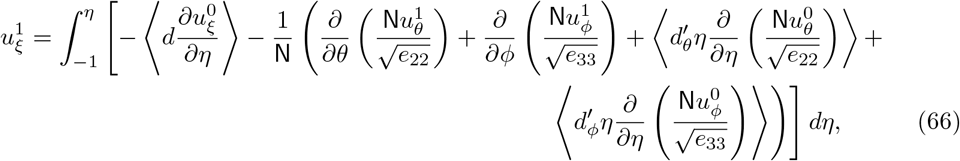

where each term can be simplified as follows

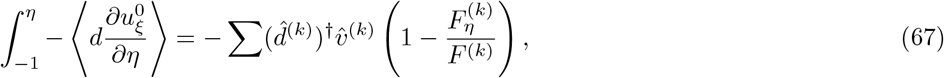

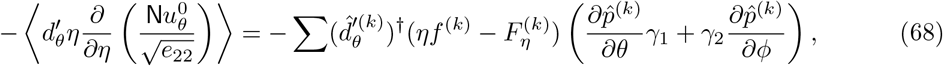

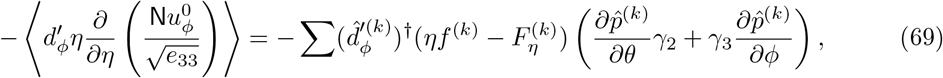

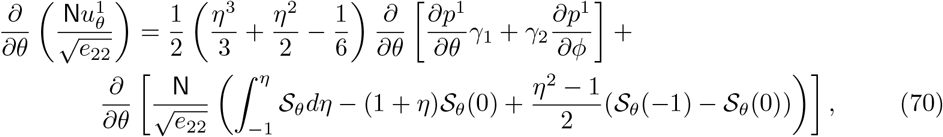

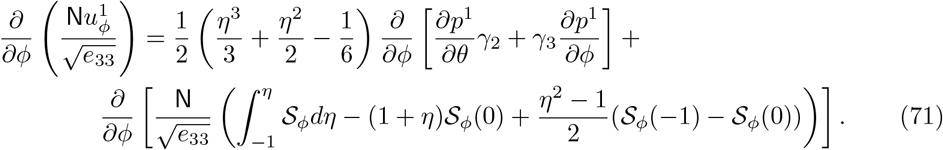

### B.3 Functions for 𝒪 (*ϵ*) analytical solution

The functions *H*_*i*_, *i* = 1, 2 from (36) can be written as 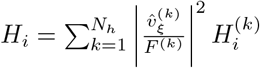. Similarly, constants *J*_*i*_, *i* = 1, 2 from (40) can be expressed as 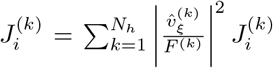. The corresponding terms in these sums are

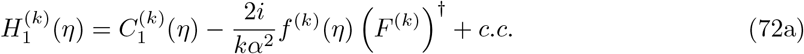

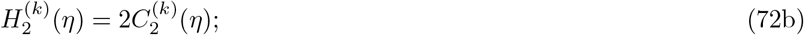

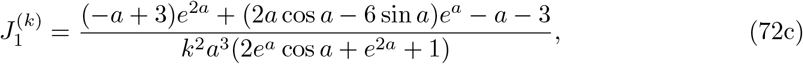

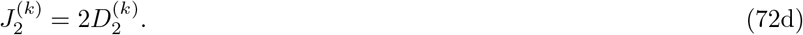

Furthermore, here we report the expression for 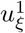,

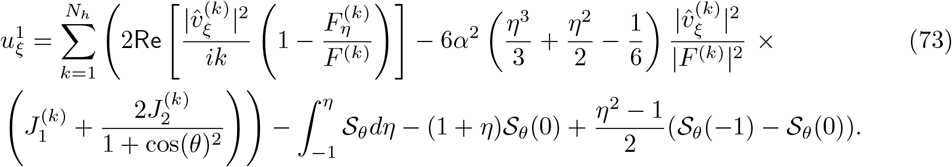

### B.4 Christoffel symbols

In this section we report Christoffel symbols, calculated as 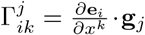 By this definition, 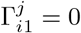, as our basis vectors do not depend on *ξ*. Other symbols appearing in inertial terms (57) are

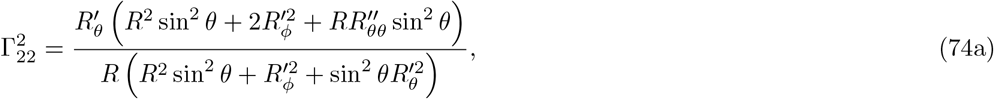

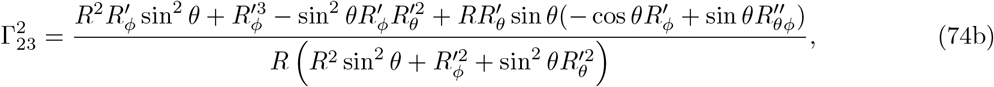

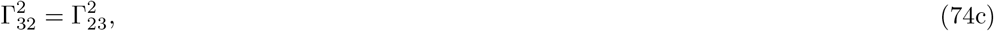

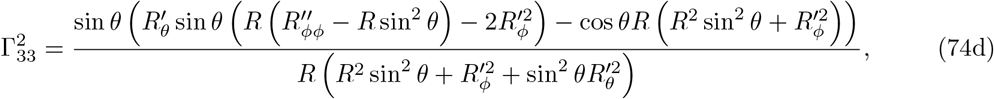

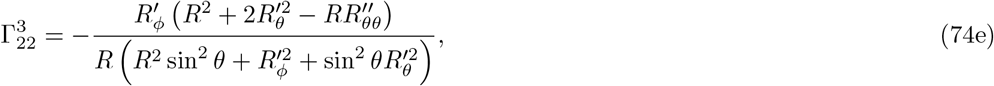

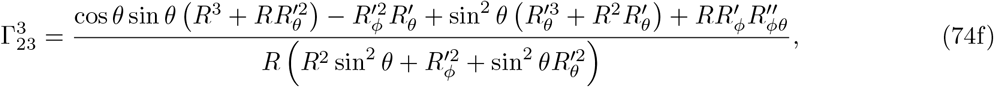

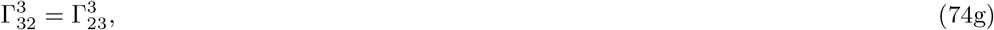

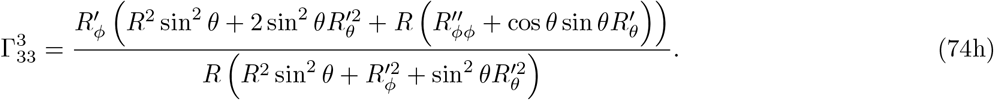

## Notes

### Competing Interest Statement

The authors have declared no competing interest.

